# De novo design of a *β*-helix tau protein scaffold: An oligomer-selective vaccine immunogen candidate for Alzheimer’s disease

**DOI:** 10.1101/2023.06.08.544248

**Authors:** Adekunle Aina, Shawn C.C. Hsueh, Ebrima Gibbs, Xubiao Peng, Neil R. Cashman, Steven S. Plotkin

**Author notes:** Current address: Center for Quantum Technology Research, School of Physics, Beijing Institute of Technology, Beijing, China 100081 (Xubiao Peng).

## Abstract

Tau pathology is associated with many neurodegenetive disorders, including Alzheimer’s disease (AD), where the spatio-temporal pattern of tau neurofibrillary tangles strongly correlates with disease progression, which motivates therapeutics selective for misfolded tau. Here, we introduce a new avidity-enhanced, multi-epitope approach for protein misfolding immunogen design, which is predicted to mimic the conformational state of an exposed epitope in toxic tau oligomers. A predicted oligomer-selective tau epitope ^343^KLDFK^347^ was scaffolded by designing a *β*-helix structure that incorporated multiple instances of the 16-residue tau fragment ^339^VKSEKLDFKDRVQSKI^354^. Largescale conformational ensemble analyses involving Jensen-Shannon Divergence and the embedding depth 𝒟 showed that the multi-epitope scaffolding approach, employed in designing the *β*-helix scaffold, was predicted to better discriminate toxic tau oligomers than other “monovalent” strategies utilizing a single instance of an epitope for vaccine immunogen design. Using Rosetta, 10,000 sequences were designed and screened for the linker portions of the *β*-helix scaffold, along with a C-terminal stabilizing *α*-helix that interacts with the linkers, to optimize the folded structure and stability of the scaffold. Structures were ranked by energy, and the lowest 1% (82 unique sequences) were verified using AlphaFold. Several selection criteria involving AlphaFold are implemented to obtain a lead designed sequence. The structure was further predicted to have free energetic stability by using Hamiltonian replica exchange molecular dynamics (MD) simulations. The synthesized *β*-helix scaffold showed direct binding in surface plasmon resonance (SPR) experiments to several antibodies that were raised to the structured epitope using a designed cyclic peptide. Moreover the strength of binding of these antibodies to *in vitro* tau oligomers correlated with the strength of binding to the *β*-helix construct, suggesting that the construct presents an oligomer-like conformation and may thus constitute an effective oligomer-selective immunogen.

## Introduction

Tau is a microtubule-associated protein mainly found in the axonal part of neurons (*1*). The assembly of *α* and *β* tubulins results in the formation of microtubules. Tau binding to tubulin promotes the assembly of microtubules and stabilization of the microtubule network, which is required for axonal transport and neuronal health (*2 –4*). A hallmark of pathology in Alzheimer’s disease as well as other tauopathies is the formation of tau oligomers and fibrils (*5 –6*), which in turn destabilizes microtubules and thus intracellular trafficking (*7*). Destabilization of microtubules leads to neuronal and synaptic loss (*8 –9*), and promotes neuroinflammation (*10 –11*). Neurodegenerative disorders involving tau pathology, collectively known as tauopathies, include frontotemporal dementia (FTD) or Pick’s disease (PiD), chronic traumatic encephalopathy (CTE), corticobasal degeneration (CBD), progressive supranuclear palsy (PSP), and Alzheimer’s disease (AD) (*12*). The presence of tau lesions strongly correlates with the progression of these diseases (*10, 13 –15*). Alternative splicing gives rise to alternate isoforms having variable numbers of microtubule binding domains (*12*). During pathology, a given isoform may aggregate on a different energy landscape (*16*) to various fibril morphologies that correspond to different disease phenotypes (*17 –19*).

While the pathogenesis is not yet fully understood, the aberrant presence of extracellular Amyloid-*β* (A*β*) plaques and intracellular tau-containing neurofibrillary tangles in AD brains make both A*β* and tau proteins the two main therapeutic targets (*6*). Safely targeting therapeutically relevant epitopes on A*β* or tau is nevertheless a herculean task (*20*). Immunogens (or vaccines) are highly effective therapeutic agents employed in the public health sector to trounce diseases. After two decades of research and development, vaccines for treating AD have been unsuccessful (*21 –24*). However, the recent approval of the first immunotherapy against AD by the United States Food and Drug Administration (FDA), although amidst some controversy, has reactivated interest in immunotherapeutic strategies for combating AD. AADvac1, the first immunotherapy targeting tau, was advanced to clinical trials in 2013 (*25*). Over the following decade, more than a dozen immunogens targeting tau have been developed and advanced to clinical trials (*10*). A strategy employed in some administrations of peptide or protein immunogens in neurodegenerative disease is the presentation of either whole protein or a peptide containing an epitope of interest. Nevertheless, it is the aggregated, oligomeric forms of the protein which have been shown to be toxic (*6*). This motivates a strategy wherein key epitopes of the protein are scaffolded in such a way as to present multiple copies of the epitope in a polyvalent immunogen.

The problem of designing an effective vaccine to treat a protein misfolding disease such as AD is two-fold, involving both epitope identification and selective targeting to pathogenic species. Epitopes must be identified that are disease-specific and induce strong and longlasting immunity (*26*), and the presentation of the immunogenic, disease-specific epitopes must selectively target pathogenic species while sparing healthy proteins. The epitope in the healthy protein often has an identical and unmodified primary amino acid sequence as the toxic species. Vaccines induce an ensemble of immunogen-specific antibodies, which recognize various epitopes on the immunogen. T cells recognize MHC-presented linear peptide sequences, while B cells recognize both linear and conformational epitopes that are exposed on antigenic surfaces (*24 –27*). In the present application, an effective vaccine must induce antibodies that selectively bind to misfolded conformations in toxic forms of the protein over healthy “native” conformations (*28*). Like its predecessors tested in clinical trials, there remain concerns regarding the efficacy of the recently approved A*β*-targeting antibody aducanumab for the treatment of AD (*29 –32*). It is therefore important to continue to innovate new approaches to combat AD.

In this paper we rationally design an immunogen that captures the conformational ensemble of misfolding-specific epitopes in tau protein, characterized by regions most readily disordered and solvent-exposed in a stressed, partially disordered fibril. We achieved this by designing a scaffold that is primarily a *β*-helix that incorporates multiple instances of the targeted epitope, and which is stabilized by a single *α*-helix. Conformational analyses show that this scaffold presents the misfolding-specific epitopes in conformations distinct from those in either the intrinsically disordered monomer, or the amyloid fibril, more effectively than the approach of scaffolding a single epitope. The designed immunogen is predicted to be suitable for eliciting therapeutic antibodies that are selective for toxic tau oligomers in Alzheimer’s disease.

## Results and Discussion

### Prediction of tau epitopes

We predicted misfolding-specific epitopes in tau protein using the procedure described in Prediction of epitopes section, which has previously been used to predict misfolding-specific epitopes in *α*-synuclein (*28 –33*), superoxide dismutase (SOD1) (*34*), and A*β* (*35 –36*). An equally weighted sum of three measures of local disorder from the fibril structure (PDB 5O3L; see Fig. S1), including loss of native contacts (ΔQ), increased root mean squared fluctuations (ΔRMSF), and increased solvent accessible surface area (ΔSASA) were used to predict epitopes (Fig.s S2, S3, S4). Fig. 1A shows the per residue normalized values for ΔSASA, ΔQ, and ΔRMSF. As necessary condition for a candidate epitope, a contiguous string of residues must satisfy a consistent decrease in Q, an increase in SASA, and a value of ΔRMSF greater than the average across all residues in 8 of 10 independent simulations for all metrics. Figure 1B shows the per residue weighted sum of ΔSASA, ΔQ and ΔRMSF over all chains with the boundary chains A, B, I, and J in the fibril structure having half the weight of other chains. Mathematically, we ploted the function Ω (Eq. 1) versus the residue index.

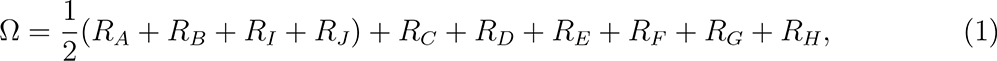

**Figure 1:**
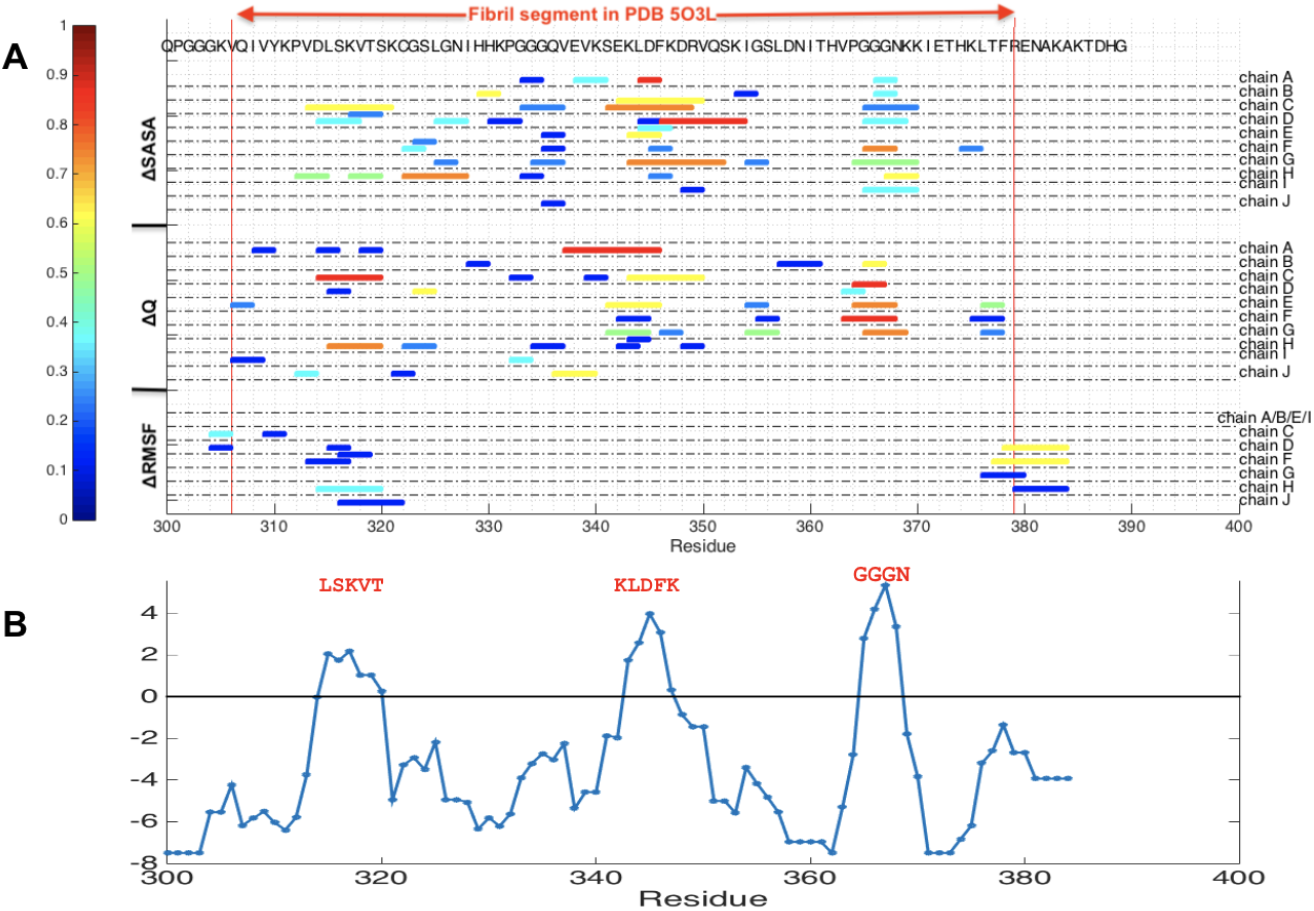
Prediction of tau epitopes. (**A**) Per residue normalized increase in solvent accessible surface area (ΔSASA), loss of native contacts (ΔQ), and local increase in root mean squared fluctuations (ΔRMSF). **(B)** Per residue weighted sum of ΔSASA, ΔQ and ΔRMSF (*28 –34*) over all chains with the boundary chains in the 5×2-chain, 2-protofibril structure A, B, I, and J having half the weight of other chains. The zero point was set to a numerical value (7.5) that identified epitopes that are within 5 amino acid long. The sequences shown in red font are the predicted epitopes for tau.

Where

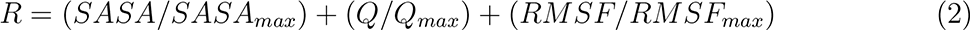

The zero point of function Ω was set at 7.5 to identify epitopes that are about 5-residue long. Three motifs above the zero point, ^315^LSKVT^319^, ^343^KLDFK^347^, and ^365^GGGN^368^, were thus predicted as misfolding-specific epitopes for tau.

#### Robustness of epitopes

The disorder-prone epitope, KLDFK, was predicted solely by analyzing the unfolding trajectories of a single tau fibril structure (PDB 5O3L). While a more robust measure would consider collective coordinates prediction for other tau fibril structures in the Protein Data Bank (PDB), such analysis would currently require a significant amount of computational resources. On the other hand, the disorder-prone epitope, KLDFK, is found to be at least moderately exposed in all of the five tau fibrils we examined, including 5O3L (*17*), 5O3T (*17*), 7P66 (*19*), 7P68 (*19*), 6TJO (*37*) (Fig. S7). We note that these fibril structures correspond to different tauopathies (Alzheimer’s, Globular Glial Tauopathy, Corticobasal degeneration) and need not expose the same epitopes. Examination of the SASA profiles in Fig. S7 shows that the 5-residue segment that containing ^343^KLDFK^347^ is partly solvent exposed, having an SASA that exceeds more than 70% of the other windows in all fibril structures analyzed, yet the epitope is generally flanked by regions having even higher solvent exposure. We may thus hypothesize that this epitope would experience a significant change in solvent exposure upon stress to the fibril. The pairwise local distance test (lddt) (*38*) shows that 5O3L and 5O3T as well as 7P66 and 7P68 are mutually similar, resulting in 3 distinct structure classes (Fig. S7 G).

#### Immunogenicity of epitopes

The immunogenicity of the epitopes predicted from Collective Coordinates (*34*) was examined using the Epitopia (*39*) server. Fig. S10 plots the immunogenicity for each residue on a scale between 1 (lowest) and 5 (highest). By this metric ^343^KLDFK^347^ had below-average immunogenicity (2.5 *vs.* 3.3 average for 5 aa segments). Nevertheless, antibodies have been raised to cyclic peptides containing KLDFK, as discussed further in Section [Results: Binding strength of antibodies…]. Interestingly, ^315^LSKVT^319^, another misfolding-prone epitope predicted from the collective coordinates algorithm, did have high predicted immunogenicity.

In summary, a protofibril region is used as a model to predict oligomer-selective epitopes. These regions of peptide sequence are conformationally distinct from either those in an isolated monomer (because they are presented in a stressed fibril conformation), or those in the fibril itself (because they were selected as being disordered from the fibril when stressed). This recipe has been previously successful in identifying oligomer-selective anti-bodies in A*β* (*35 –36*) and *α*-synuclein (*28 –33*). The problem then turns to how to properly scaffold the predicted epitope in order to present it in a conformation similar to that in the stressed fibril. For A*β* and *α*-synuclein, this has been successfully achieved by cyclic peptide “glycindel” scaffolds (*28*). We now turn to this analysis for the above-predicted epitopes.

### Conformational selectivity of single-epitope cyclic peptide scaffolds

Our goal at this stage is to design scaffolds that are selective for toxic tau oligomers. In other words, we aim to design scaffolds that present epitopes in similar conformations to those present on toxic tau oligomers, that are different from conformations presented by healthy tau monomers, or fibril, as we wish to target paracrine species that have the potential to spread pathology by prion-like propagation (*6 –40*)

To this end, we computationally designed “glycindel” cyclic peptides (see Methods: Epitope scaffolding using cyclic peptides) to scaffold two of the three predicted tau epitopes, including ^315^LSKVT^319^ and ^343^KLDFK^347^ epitopes. We did not construct scaffolds for the ^365^GGGN^368^ predicted tau epitope because it consists of mostly glycine (GLY) residues, which are less immunogenic than other side chains. Each constructed cyclic peptide scaffold consists of a single instance of either the ^315^LSKVT^319^ or ^343^KLDFK^347^ epitope flanked by between one to four variable numbers of GLY on the Nand/or the C-terminus. Hence, we constructed a total of 4×4 or 16 cyclic peptide scaffolds for each of the ^315^LSKVT^319^ and ^343^KLDFK^347^ epitopes.

To determine the conformational selectivity of the constructed cyclic peptide scaffolds *in silico*, we sampled the conformational ensemble of each epitope in four different contexts: The fibril, oligomer, monomer, and each of 16 × 2 = 32 cyclic peptide scaffolds, by performing all-atom MD simulations in explicit solvent (see Methods: Sampling conformational ensembles). Each cyclic peptide epitope ensemble was then compared to the same epitope ensemble in the context of the fibril, oligomer, and monomer. We used Jensen-Shannon Divergence (JSD) to quantify the degree of similarity between two conformational ensembles (see Methods: Comparing conformational ensembles). We also apply a metric we have recently introduced to measure the degree to which one ensemble is subsumed by another ensemble, the embedding depth 𝒟 (*28*).

Glycindel scaffolds were generically distinct from the fibril, with JSD values between scaffold and fibril ranging from 0.96 to 1.00. We thus focused on comparisons with the monomer and stressed fibril oligomer model ensembles.

We seek epitope scaffolds with both high similarity (low JSD) to the oligomer ensemble, and low similarity (high JSD) to the monomer ensemble. Similarly, we seek epitope scaffolds with both high embedding depth (high D) in the oligomer ensemble, and (low D) in the monomer ensemble. Fig. 2 A shows the conformational ensemble similarity (JSD) to the oligomer *vs.* JSD to the monomer, for each of the 32 cyclic peptide scaffolds (16 scaffolds each for tau ^315^LSKVT^319^ and ^343^KLDFK^347^ epitopes).

**Figure 2:**
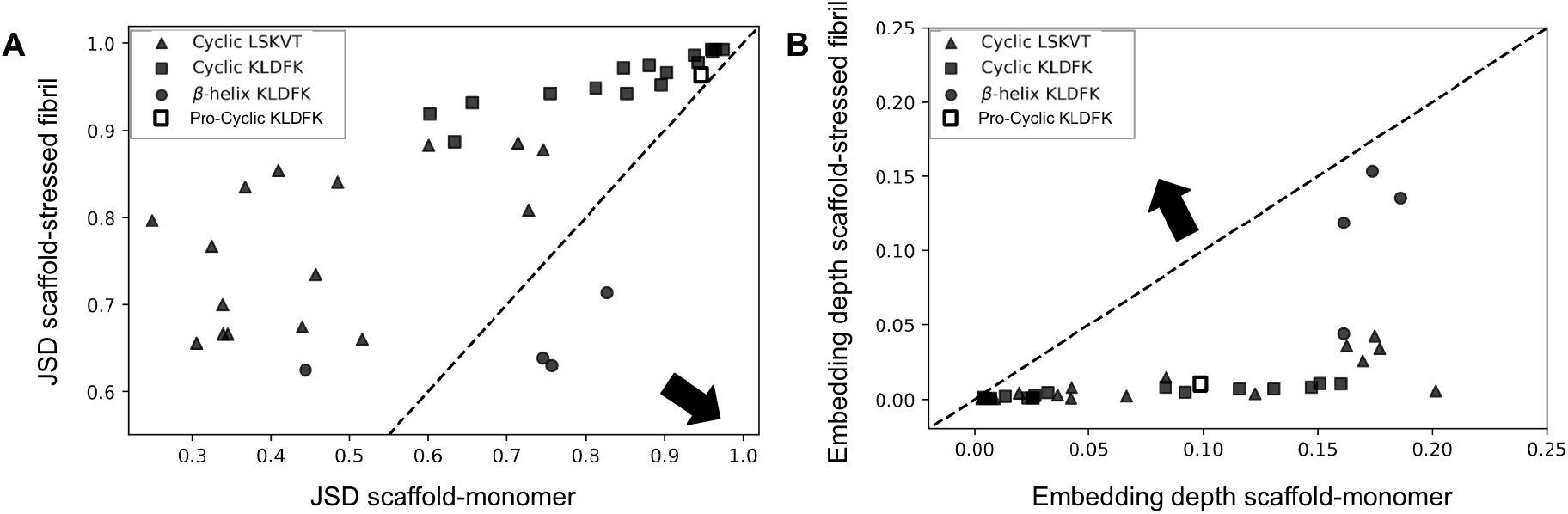
Conformational ensemble similarity. (**A**) The similarity between two conformational ensembles was quantified using JSD (see Methods: Comparing conformational ensembles), which is normalized to lie between 0 (for identical ensembles) to 1 (for entirely different ensembles). Shown are the similarity between the epitope ensemble in the context of a scaffold and the oligomer, versus the similarity between the epitope ensemble in the context of the scaffold and the monomer, for cyclic peptide “glycindel” scaffolds of the LSKVT epitope (triangles), cyclic peptide glycindel scaffolds of the KLDFK epitope (squares), CRISPro-designed cyclic CPPPPKLDFKGPGG scaffold (open squares) and the *β*-helix scaffold of the four KLDFK epitope repeats (circles). The arrow indicates the direction of favorable conformational profile and the dashed line is the line *y* = *x*. (**B**) Same as for panel A, but for the embedding depth D. Now, data points towards the upper left above the dashed *y* = *x* line are favorable; interestingly none of the points fall in this regime, but the *β*-helix scaffold epitopes do cluster differently, and have the largest embedding depth with the oligomer model. Cyclic peptide ensembles have variable embedding depth with the monomer ensemble, but they show little embedding in the oligomer model ensemble.

Several observations are apparent from Fig. 2 A. In general, ^315^LSKVT^319^ scaffolds sample epitope ensembles that are more similar to both the monomer and the oligomer ensembles than ^343^KLDFK^347^ scaffolds. There is a positive correlation between the similarity between scaffold and oligomer and the similarity between scaffold and monomer, with Pearson correlation coefficient *r* = 0.71 for all the cyclic peptide scaffolds, moderate correlation for the ^315^LSKVT^319^ epitope scaffolds (*r* = 0.56), and strong correlation for the ^343^KLDFK^347^ epitope scaffolds (*r* = 0.92). This correlation shows that as the cyclic peptide scaffolds become less similar to the monomer (increasing JSD), which is desirable, they also become less similar to the oligomer (increasing JSD), which is undesirable. It is worth noting that the JSD between monomer and oligomer is 0.54, more similar than any of the scaffolds to the oligomer, and more similar than any of the KLDFK scaffolds to the monomer. The more similar the monomer and oligomer ensembles are, the stronger the correlation we expect in the data of Fig. 2 A.

The desired scaffold should be more similar to the oligomer than the monomer, which would put the data point for a scaffold below the *y* = *x* dashed line in Fig. 2. The result here shows an unfavorable result that the typical conformations explored by glycindel cyclic peptide scaffolds of ^315^LSKVT^319^ and ^343^KLDFK^347^ epitopes are predicted to be more similar to those of an isolated monomer of tau than an oligomer model using a stressed fibril. We note that this result is in contrast to glycindel scaffolding of epitopes for *α*-synuclein, which were able to predict scaffolds having greater similarity to oligomers (*28*).

Fig. 2B shows the embedding depth 𝒟 of each of the 32 cyclic peptide ensembles in the oligomer ensemble *vs.* 𝒟 in the monomer ensemble, as well as the corresponding embedding depths for the 4 repeats of the epitope in the *β*-helix scaffold. Here, the glycindels show variable embedding depth in the monomer ensemble, but show very little embedding in the oligomer model ensemble (points are spread horizontally along the bottom of the plot, below the line *y* = *x*). The *β*-helix scaffolded epitopes also embed deeper in the monomer ensemble than the oligomer ensemble, however, the 3 most structured *β*-helix epitopes cluster together and have significantly higher embedding within the oligomer ensemble than do any of the glycindels. The N-terminal weakly structured epitope in the *β*-helix scaffold also behaves more monomer-like, falling within the distribution of the glycindel scaffolds.

### Design of avidity-enhanced multi-epitope ***β***-helix scaffolds

From the above result, all of the 32 cyclic peptide scaffolds are predicted to explore monomer-like conformations more often than oligomer-like conformations. Though this does not rule out the possibility that antibodies raised against such a peptide may be selective to the oligomer-like conformations occasionally explored by the ensemble, it suggests that the probability for such a scenario is low. We therefore designed a single-chain *β*-helical scaffold consisting of multiple instances of the ^343^KLDFK^347^ epitope. We chose the ^343^KLDFK^347^ epitope because of its location in the tau fibril structure (Fig. 6 A). This epitope is in the 4th microtubule binding domain of tau; it does not overlap with the epitopes of any antibodies currently in clinical trials for AD (*6 –41*). The closest epitope is that of Zagotenemab, which has a discontiguous epitope consisting of residues 7–9/312–322. (*42*) The epitope ^343^KLDFK^347^ is a motif which is part of a larger contiguous sequence in the tau monomer, ^339^VKSEKLDFKDRVQSKI^354^ (tau339-354), that has close spatial proximity between its Nand C-termini within the tau fibril structure PDB 5O3L. Tau structures in the PDB are polymorphic, often depending on the disease phenotype. We have chosen a tau structure associated with AD, however other tau structures in the PDB also satisfy the condition that the N- and C-termini of V339 and I354 are in close spatial proximity (e.g. PDBs 6NWP and 6NWQ involved in chronic traumatic encephalopathy). The strategy used here should potentially be applicable to this disease as well. Beta-helical structures have been observed in nature in other contexts, for example antifreeze proteins (*43 –44*).

To design *β*-helix peptide scaffolds, we tethered 5 copies of tau339-354 the sequence with glycine (GLY) tripeptides (GGG) (see Fig. 6). We then made a S352C (serine 352 to cysteine (CYS)) mutations in the first four copies of the sequence, with the intention of designing two disulfide bridges that would improve the stability of the scaffold (Fig. 3 A). Indeed AphaFold (*45*) predicts a *β*-helix structure (Fig. 3 B) with the S352C substitutions, but predicts a disordered peptide without the S352C substitutions (see Methods: Structure prediction with AlphaFold). All 5 AlphaFold models predicted a *β*-helix structure for the five-repeat sequence (with S352C substitutions) with high average per residue confidence scores of 89%, 92%, 89%, 88%, and 85% for the 1st through 5th AlphaFold models, respectively (see Fig. S5C).

**Figure 3:**
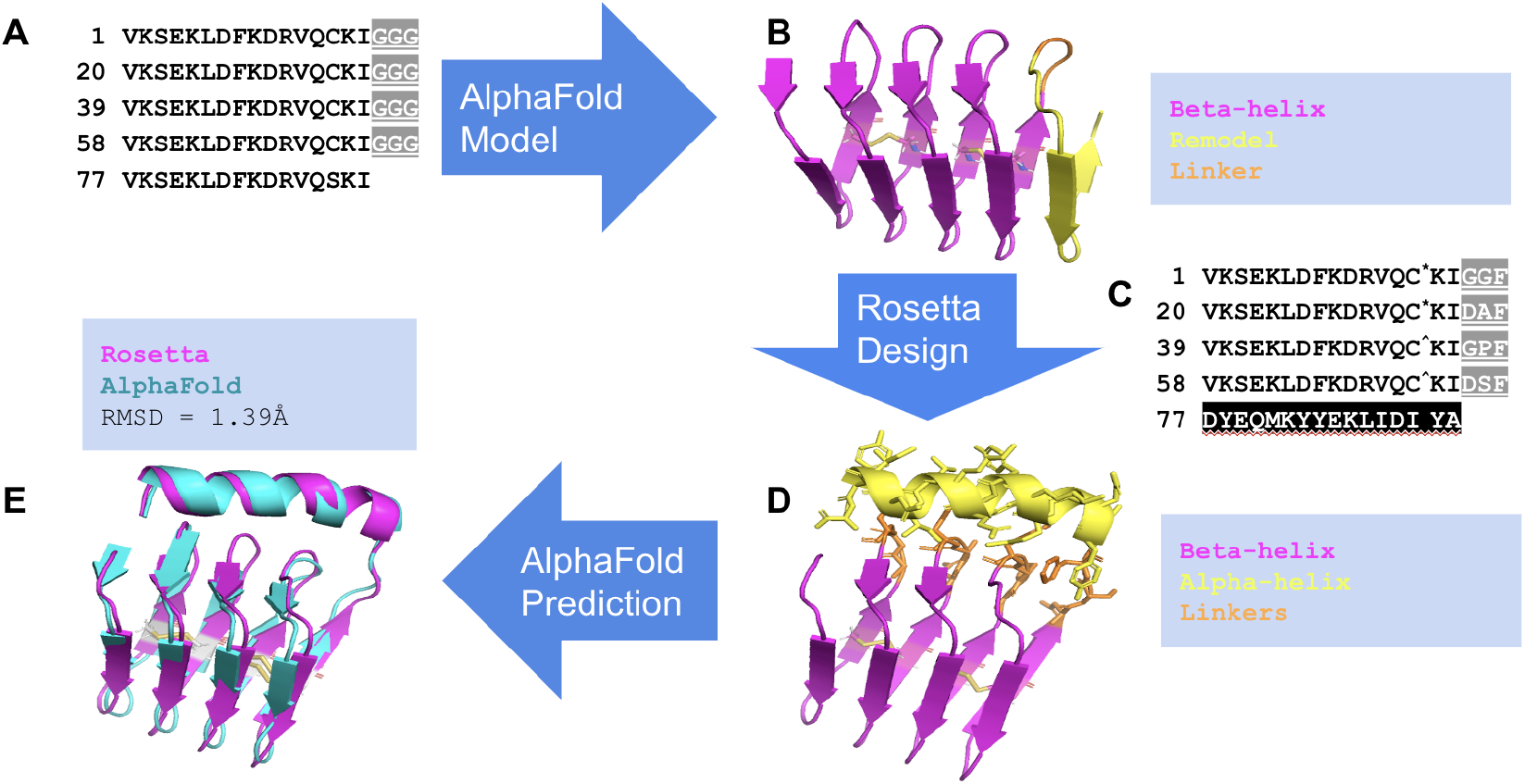
*β***-helix scaffold design strategy.** (**A**) Five copies of tau-derived 16-residue ^339^VKSEKLDFKDRVQSKI^354^ repeating sequence are linked serially by glycine tripeptide linkers (GGG); S352C substitutions are made in the first four repeats. These were used to model (**B**) a *β*-helix structure, with disulfide bridges between residues C14 and C33 and residues C52 and C71. (**C**) Redesign of the last repeat (black box) and linker residues (gray boxes) to improve the stability of (**D**) the *β*-helix scaffold by creating interactions between the designed residues. Last repeat is shown in yellow and linkers are shown in orange, both with licorice side chains. (**E**) The designed *β*-helix structure (magenta) and AlphaFold predicted structure (cyan) have C*_α_* RMSD of 1.39Å. All images were rendered with PyMOL molecular visualization system (https://pymol.org).

We had also examined a 4-repeat *β*-helix scaffold (Fig. S6 A). Although the AlphaFold confidence (PLDDT) scores for this construct were high (Fig. S6 B), the free energy surface as a function of *Q*, obtained by Hamiltonian replica exchange molecular dynamics (HREMD) simulations, did not predict a stable native-like structure (Fig. S6 C, see also Results: The designed *β*-helix scaffold is structurally and energetically stable). This prompted additional design as described further below.

The formation of *β*-helix structure always put cysteines C352 in successive repeats of tau339-354 within 4.8 Å of each other. Two disulfide bonds, between cysteines in the 1^st^and 2nd repeats, and between cysteines in the 3rd and 4th repeats, were predicted as intended by only the third and fifth AlphaFold models; the other three models predicted only one of the two disulfide bonds.

We thus used the 3rd AlphaFold model with the two designed disulfide bonds and higher confidence score (89%) as the initial *β*-helix structural model for redesign. To further improve the stability of *β*-helix structural model, we remodeled the amino acids starting from the last GGG linker and including the linker and the last tau339-354 repeat (see Methods: *β*-helix structure modeling). The last tau339-354 repeat was remodeled to an *α*-helix structure in a conformation that was docking and interacting with all the linkers (Fig. 3 D). For both linker and *α*-helix, we allowed for the possibility of any of the 20 naturally occurring amino acids to occur in all remodeled positions. Finally, all four linkers and *α*-helix residues were designed to optimize the sequence and structure of the *β*-helix scaffold in (Fig. 3 D) (see Methods: *β*-helix sequence design and structure optimization).

Ten thousand (10,000) decoys of *β*-helix scaffold were generated during the design phase using Rosetta (*46 –47*). Fig. S5 A plots the Rosetta energy per residue of the designed decoys against the C*_α_* root mean squared deviation (RMSD) from the starting Rosetta-remodeled structure, with energy = 1.98 Rosetta energy units (REU). The RMSD for all designed decoys is below 1 Å except for one outlying decoy with an RMSD of 1.13 Å. All the designed decoys have energies below -3 REU per residue, which is significantly lower than the energy of the starting structure.

### AlphaFold predicts the designed ***β***-helix structure

To validate the feasibility that a designed sequence folds into the designed *β*-helix structure, AlphaFold was used to predict the structure from the designed sequences alone (see Methods: Structure prediction with AlphaFold). A distribution of energy was constructed for the 10,000 decoys and the lowest 1% of energy defined a cutoff that included 82 unique sequences. The structures of these 82 unique sequences were then predicted with AlphaFold. AlphaFold outputs 5 models per sequence, with each model predicting one structure. Table S1 shows the average per residue confidence (PLDDT) score for the median ranking model of the 5, and the C*_α_* RMSD from the Rosetta-designed *β*-helix structure for the three top ranking models for 28 selected sequences. The 28 selected sequences are those from the 82 lowest energy sequences that satisfy two criteria: 1.) They have an average per residue confidence score for the median ranking model ≥ 50%, and 2.) each of the three top ranking models has a C*_α_* RMSD from the remodeled *β*-helix structure ≤ 5 Å.

To characterize the structural and energetic stability of a representative designed sequence *in silico*, several selection criteria were imposed. We selected a sequence that 1) has a low energy; it is among the 1% lowest energy decoys, corresponding to an energy cutoff of −3.41 REU per residue, 2) has a high structure prediction score; a median confidence score (across 5 models) ≥ 70, which is the threshold for high confidence structure prediction for AlphaFold (*45*). That is, at least three AlphaFold models must have high confidence scores (see Fig. S5 B), and 3) has the designed structure across all the top three models; it is predicted to have i) the designed *β*-helix structure, and ii) correctly forms the two designed disulfide bonds. Table S1 shows that model 0 is the only sequence satisfying this stringent selection criteria. The Rosetta designed structure and AlphaFold predicted structure for model 0 have C*_α_* RMSD of 1.39Å (Fig. 3 E).

### The designed ***β***-helix scaffold is structurally and energetically stable

One of the most important pharmacological properties for a peptide such as the designed tau *β*-helix scaffold intended for use as an immunogen is stability. To this end, the stability of the selected *β*-helix scaffold (i.e. model 0) was characterized *in silico*. All simulations were performed in explicit solvent (0.15mM NaCl) at 300K and 1bar using GROMACS (*48*) with the CHARMM36m force field (*49*), and periodic boundary conditions.

#### Metastability

To check the structural stability of the scaffold in the free energy basin of the designed structure, a 100 ns all-atom MD simulation starting from the designed structure was performed, and the last 50 ns was used for analysis. Heavy-atom root mean square deviation (RMSD) and C*_α_* root mean square fluctuation (RMSF) were used to quantify the global and local stability, respectively. The RMSD *vs.* time plot in Fig. 4A shows that the scaffold is globally structurally stable within the simulation time. The RMSD from the starting structure of the sampled conformations remains below ≈ 3 Å. Apart from five N-terminus residues with RMSF between 1.5 Å to 3.5 Å (Fig. 4 B), the per residue local fluctuations remain below the size of a carbon atom (1.4 Å) for all residues, suggesting a high structural stability for the scaffold in the free energy basin of the designed structure.

**Figure 4:**
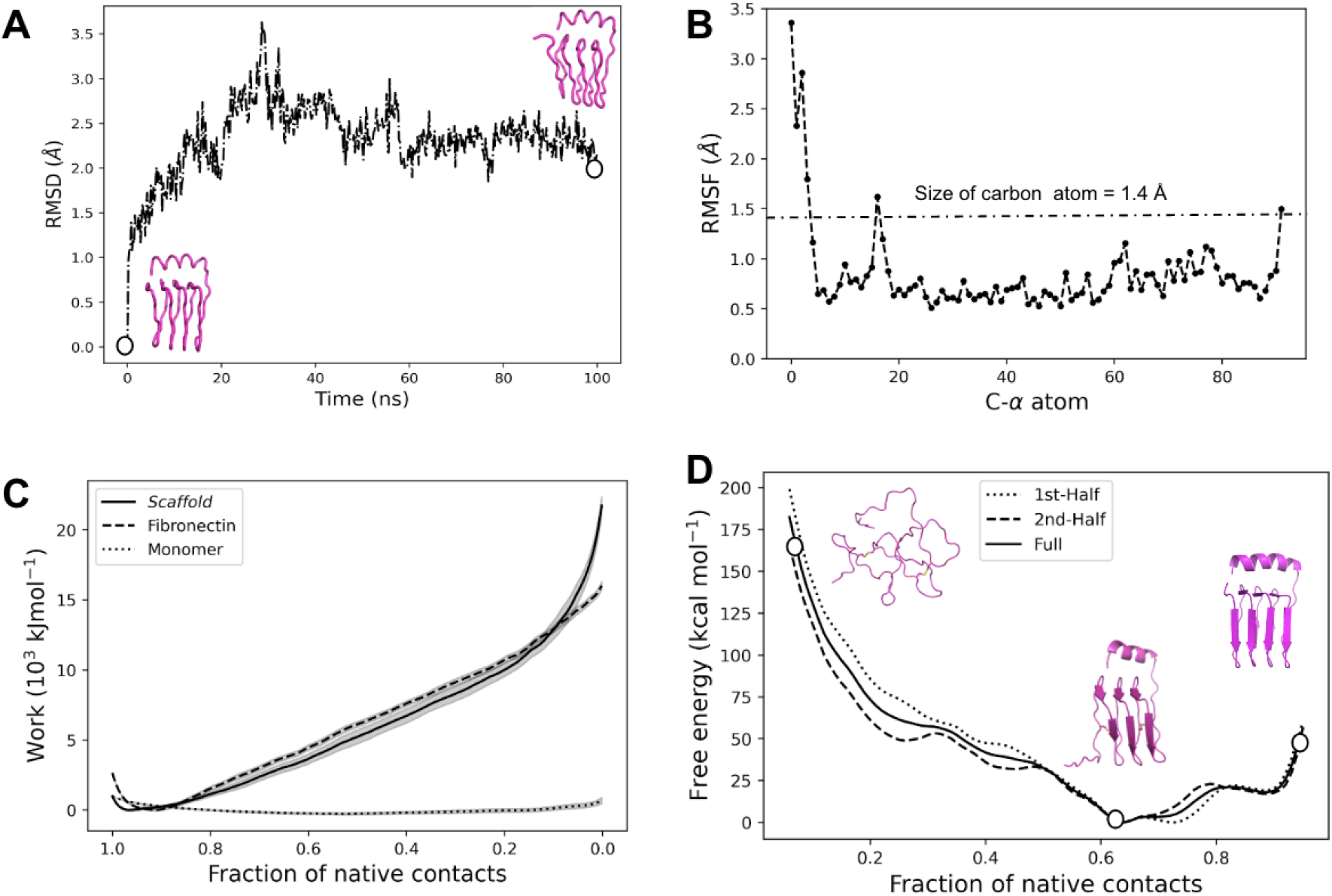
Computational characterization of the stability of *β*-helix scaffold. The (**A**) global and (**B**) local structural stability in the free energy basin of the Rosetta designed structure. Inset images in (**A**) show snapshots of the initial and final structures. The secondary structure evolution over the 100 ns simulation time is shown in Fig. S11. (**C**) The mechanical stability measured by the average cumulative work done during forced unfolding simulations (solid line), compared to fibronectin (*50 –51*), a stable extra-cellular matrix protein with similar length (dashed line), and an isolated (intrinsically disordered) tau monomer (dotted line), which serves as a negative control. (**D**) Thermodynamic stability determined from the relative free energy surface along the fraction of native contacts *Q* (Eq. 6) order parameter. Snapshots from the first half (dotted), second half (dashed), and full 80ns trajectories (solid) in each umbrella are shown (see text). Inset images show representative snapshots of the structures at the free energy minimum, where *Q* = 0.64, and where *Q* = 1 and *Q* = 0.1.

#### Mechanical stability

To quantify mechanical stability, ten independent forced unfolding simulations were performed (see Methods: Mechanical unfolding simulations). The cumulative average work done by the applied forces to unfold the *β*-helix scaffold structure as a function of the native contact order parameter *Q* (Eq. 6) was taken as a measure of mechanical stability. The work performed in a given trajectory generally averages out the stochastic effects of forces and is a smooth function of the order parameter (*52 –53*). Here, the work performed was averaged over ten independent simulations to further average out stochastic effects during the unfolding process. For comparison, the analogous calculation was performed to assess the mechanical stability of isolated fibronectin type III domain of human tenascin (PDB ID: 1TEN) (*50 –51*) and tau monomer (chain A from PDB ID: 5O3L) (*17*), which serve as positive and negative controls respectively. The force-field and simulation model (all-atom in TIP3P solvent) used for all proteins are the same, so no additional normalization across time, energy, or force is required for accurate comparison (*54*). Figure 4 C shows that the mechanical stability of the scaffold is comparable to that of fibronectin, a stable extra-cellular matrix protein with similar length (92 versus 90 amino acids).

#### Thermodynamic stability

We also determined the thermodynamic stability of the *β*-helix scaffold, by performing a Hamiltonian replica exchange molecular dynamics (HREMD) simulation to determine the free energy surface as a function of *Q* (see Methods: Hamiltonian replica exchange molecular dynamics). The HREMD simulation consists of 100 replicas with harmonic restraints centered at positions equally spaced between the fraction of native contacts *Q* = 0 to *Q* = 1. Each replica was simulated for 250 ns for a total of 25 *µ*s. A total of 8000 conformations were sampled at 10 ps intervals from the last 80 ns of each replica to determine the relative free energy of unfolding along Q for the scaffold. Figure 4 𝒟 shows a largely downhill free energy surface towards higher degrees of nativeness, until the free energy minimum of the scaffold at *Q* = 0.64. There is a weak metastable minimum near *Q* ≈ 0.9, but entropic forces bias the global minimum towards a lower degree of native contacts. Nevertheless, the global minimum ensemble has a large amount of native structural similarity:

The average RMSD of this ensemble is 1.25 Å, and if we ignore the largely disordered N-terminal 13 residues, the RMSD of the remainder is 0.45 Å. Fig. 4 𝒟 also shows representative snapshots of the conformations sampled at *Q* = 0.64, *Q* = 0.1, and *Q* = 1. The *Q* = 0.64 snapshot shows that most of the lost contacts are the result of the flexible N-terminus (c.f. Fig. 4 B). Overall, this result shows that the designed scaffold is thermodynamically stable in a largely structured, *β*-helical conformation with a stabilizing *α*-helix. This result is in contrast to similar analyses applied to a 4-repeat *β*-helix construct (without a stabilizing *α*-helix), which does not exhibit a stable native-like structure (Fig. S6 C). We note that as an immunogen proxy for an oligomer, which generally are conformationally labile and thus elusive to structural determination (*6*), some degree of partial disorder may actually be a favorable feature of the designed scaffold.

#### Proteolytic stability

Cleavage analysis by the Procleave (*55*) server indicates that the designed *β*-helix structure was predicted to have cellular stability comparable or better than ubiquitin (Fig. S8). Specifically, the highest-scoring cleavage site for the *β*-helix protein scored higher than ubiquitin for only 8 out of the 27 proteases examined.

#### Aggregation propensity

The aggregation propensity of the designed *β*-helix scaffold structure, as determined by the AGGRESCAN3D server(*56*), was found to be comparable to ubiquitin, and less than that of transthyretin, an amyloid-prone protein (*57*). Fig. S9 shows plots of the aggregation score vs. residue index, along with the mean aggregation score and the fraction of residues that have an aggregation score larger than zero. The overall aggregation propensity of the construct is observed to be intermediate to ubiquitin and transthyretin, and closer to ubiquitin. This may not be surprising, since the region of tau sequence selected for scaffolding was the first to disorder upon the simulated application of stress to the fibril. Consistent with this, the region containing this epitope is the least aggregation-prone according to AGGRESCAN3D (Fig. S9 E).

### The designed ***β***-helix scaffold is conformationally selective for a stressed protofibril model of tau oligomers

To determine the conformational selectivity of the designed *β*-helix scaffold for a stressed fibril model of tau oligomers (c.f. Fig. 2A), the conformation of the predicted ^343^KLDFK^347^ epitope was sampled in the context of monomer, stressed protofibril oligomer, and *β*-helix scaffold (see Methods: Sampling conformational ensembles). Four different KLDFK ensembles, corresponding to the four KLDFK instances in the *β*-helix scaffold sequence, were separately analyzed for the *β*-helix scaffold. Each KLDFK scaffold ensemble was then compared with the monomer and the oligomer ensembles, respectively, using the Jensen-Shannon divergence (JSD) similarity measure (see Methods: Comparing conformational ensembles), and the embedding depth 𝒟.

Here, we compare the conformational selectivity of a multi-epitope *β*-helix scaffold to single-epitope cyclic peptide scaffolds for the tau ^343^KLDFK^347^ epitope. Figure 2A shows that the *β*-helix scaffold is predicted to be more selective for tau oligomers than are cyclic peptide scaffolds. For the same conformational similarity to the oligomer, three of the four *β*-helix ensembles clearly show lower similarity (higher JSD) to the monomer than KLDFK cyclic peptide scaffolds. Similarly, for the same conformational disimilarity to the monomer, three *β*-helix ensembles show higher similarity (lower JSD) to the oligomer than all of the KLDFK cyclic peptide scaffolds. The N-terminal KLDFK motif has an ensemble that is more similar to the monomer than that of the other three epitope repeats. This is expected since the N-terminus of the *β*-helix scaffold is relatively flexible, assuming the accuracy of the force field used in the simulations (CHARMM36m (*49*)) (Fig. 4 D).

Previously, Gibbs *et al.* have constructed single-epitope cyclic peptide glycindel scaffolds to successfully generate A*β* oligomer-selective antibodies (*36*), and *α*-synuclein oligomer-selective antibodies (*33*), following a similar procedure to the one described in Epitope scaffolding using cyclic peptides section. Therefore, this analysis cannot rule out the ability of a single-epitope cyclic peptide scaffold to raise oligomer-selective antibodies, and in fact glycindel scaffolds (*28*) of collective coordinate-predicted epitopes (*34*) have been used to raise antibodies showing greater binding response to tau oligomers than tau monomers in SPR assays (*58*). Specifically, antibodies raised from glycindel scaffolds of the epitope ^343^KLDF^346^ showed greater selectivity to oligomers/monomers than a pan-tau control antibody (in SPR binding response units, RU*_olig_*/RU*_mon_* ≈ 2 on average and up to a ratio of ≈ 4 for some antibodies, while the pan-tau antibody had RU*_olig_*/RU*_mon_* ≈ 1.4). We note that this is not a huge improvement in selectivity, and the *β*-helical scaffold may result in even further enhanced selectivity. The selectivity of glycindels did show scaffold dependence (see Methods: Epitope scaffolding using cyclic peptides), e.g. antibodies raised by (3,2)KLDF were more selective to oligomers than those raised by either (3,1)KLDF or (4,1)KLDF. As well, co-incubation with antibodies raised against (3,2)KLDF showed the largest reduction of the 3 scaffolds in seeding of aggregation by PFFs in Tau RD P301S FRET Biosensor HEK cells (*58*). Taken together, this evidence suggests that conformational presentation of the epitope is important in determining the selectivity and effectiveness of the resulting antibodies. In light of the predictive data in Fig. 2, this stresses the potential fruitfulness of pursuing more accurate scaffolding strategies such as the one we have proposed here.

### Binding strength of antibodies to the ***β***-helix immunogen correlates with their binding to *in-vitro* oligomers

Several monoclonal antibodies have been obtained previously from mice immunized with the Tau cyclic peptide sequence CPPPPKLDFKGPGG (*58*). These include monoclonal antibodies 4E4, 3E2, 10B10, 6H3, 7H6, 10D4, 7E5, and 2C6, which were investigated here. An isotype negative control (mIgG1) was also used for comparison.

Although these antibodies were not raised using the *β*-helix construct, they were raised to a cyclized conformational epitope that overlaps with the sequence of the epitope in the *β*-helix construct (KLDFK), and some of these antibodies showed binding to in vitro tau oligomers (Fig. 5A). The above cyclic peptide sequence was determined from the CRISPro computational method, as described in the Methods. The ensemble of this specific sequence was not explicitly investigated, and although Fig. 2 implies that is likely distinct from the ensembles of the *β*-helix construct, the ensembles may still partially overlap, suggesting some of these antibodies may be cross-reactive.

**Figure 5:**
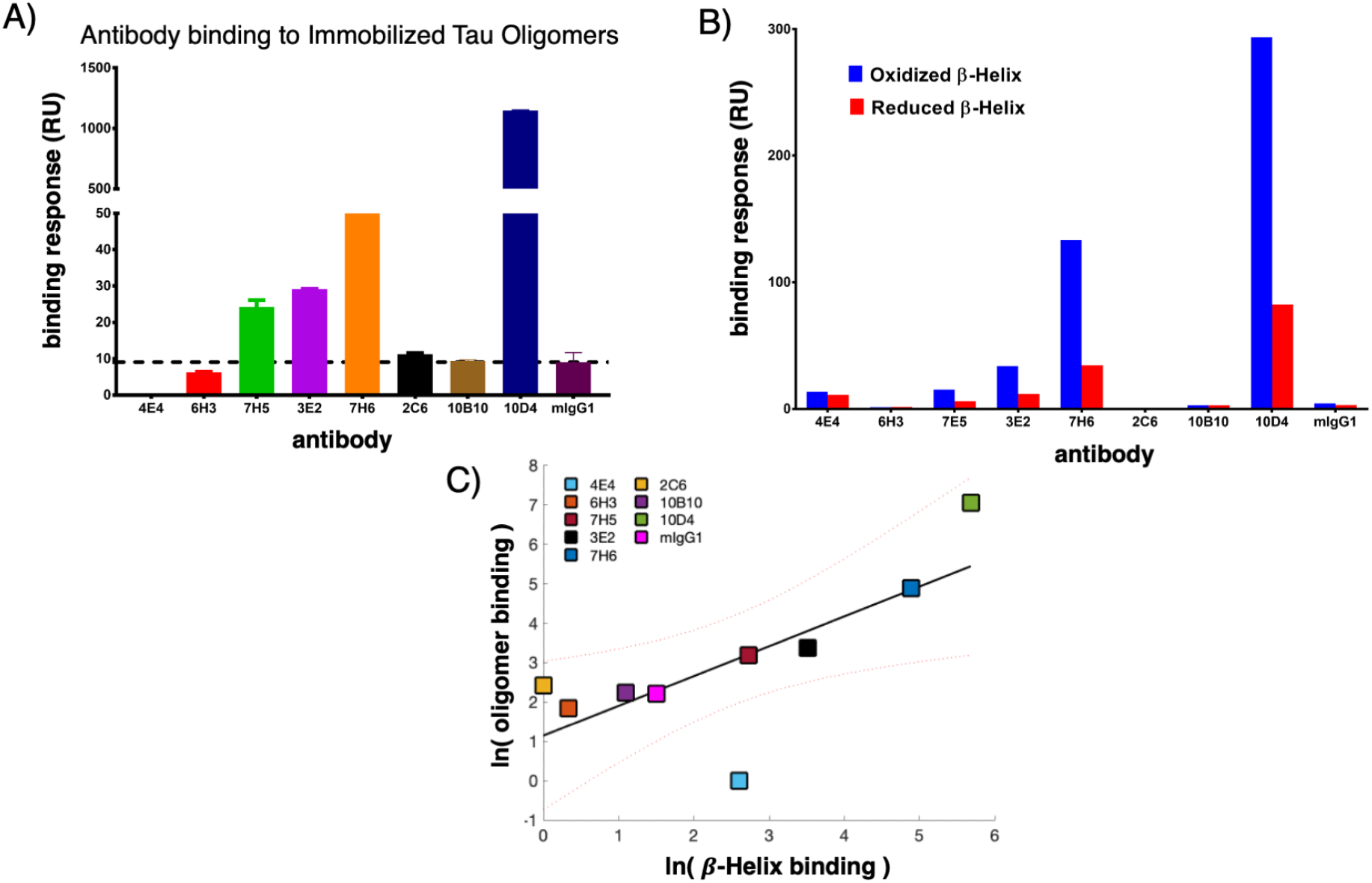
Experimental binding of antibodies to. *β***-helix construct and tau oligomers.** The (**A**) Antibodies raised to tau cyclic peptide show variable binding response (BRU) to *in vitro* tau oligomers. (**B**) Disulfide reduction of the *β*-helix construct to generate a more unfolded form resulted in decreased antibody binding, indicating that the epitope’s conformation in the *β*-helix favors antibody binding. (**C**) The strength of antibody binding to oligomers correlates significantly with the strength of binding to the *β*-helix construct (r=0.74, p=0.02), suggesting that the *β*-helix construct may present an oligomer-like conformational epitope and be a useful oligomer-selective immunogen.

Experiments to examine the conformational specificity of these antibodies to the *β*-helix construct, *vs.* an unfolded monomeric form, were performed by comparing binding of the above set of antibodies to both oxidized and reduced *β*-helix constructs. The reduced construct was used as a proxy for a more unstable, unfolded form of the construct. Antibodies raised to the epitope in the cyclic peptide ensemble were observed to bind stronger to oxidized (well-folded) *β*-helix protein, than to the reduced (more poorly folded) protein (Fig. 5B), for all antibodies tested. This result suggests that the targeted epitope is at least partially structured, and loss of structure reduces binding.

Figure 5C shows that the strength of antibody binding to oligomers correlated significantly with the strength of binding to the *β*-helix construct (r=0.74, p=0.02). This suggests that the *β*-helix construct presents an oligomer-like conformation, and thus that the *β*-helix construct may be a useful oligomer-selective immunogen.

## Methods

### Prediction of epitopes

To predict tau epitopes, two conformational ensembles were generated, one for tau fibril and the other for a partially unfolded fibril, which serves as a model for tau oligomers. This oligomer-selective epitope prediction method posits that a partially unfolded protofibril ensemble is enriched in oligomer-selective conformational epitopes (*34*), and has been supported by previous experimental evidence (*35 –36*).

The fibril ensemble was generated by performing a 30.1 ns molecular dynamics (MD) simulation, starting from a tau fibril structure (PDB ID: 5O3L) extracted from an AD patient brain (*17*). To generate the partially unfolded fibril ensemble, ten independent 150 ns bias MD simulations were performed in order to average over the stochastic process of unfolding. During the first 50 ns of the bias simulations, the number of native contacts was linearly decreased to 60% of its native value. A native contact was defined being formed when heavy atoms belonging to residues with a sequence separation ≥ 3 are within 4.8 Å over 95 percent of the time in a 100 ns equilibrium fibril simulation. This determines the total number *N* of native contacts. After 50 ns, the bias of 60% native contacts was fixed for the remaining 100 ns of simulation.

The bias potential *V* (*Q, t*) is implemented to globally unfold the tau fibril structure along the fraction of native contacts *Q* reaction coordinate, which takes the following form:

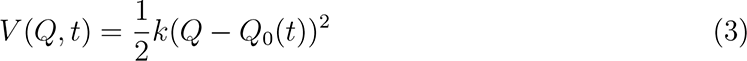

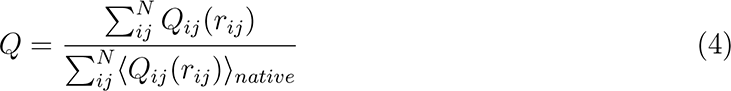

In Eq. (3), the sum in the numerator runs over the *N* native contacts that may be formed in a given conformation, while the sum in the denominator runs over the *N* native contacts in a member of the native conformational ensemble and is then averaged over conformations sampled during the 100 ns equilibrium simulation of the native fibril structure. *Q_ij_* in Eq. (4) is defined by a contact function for each atom pair *i, j* with contact separated by distance *r_ij_*, using the following formula:

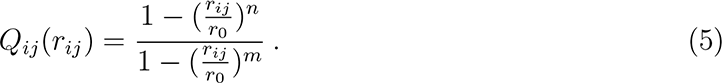

In Eq. (5), we take *r*_0_ = 4.8 Å, *n* = 6 and *m* = 12, to obtain a sigmoidal function that rapidly goes to one as *r_ij_* falls below *r*_0_, and rapidly goes to zero as *r_ij_* becomes larger than *r*_0_. All simulations were performed in explicit solvent (TIP3P water model, and 150 mM NaCl) at 300 K and 1 bar using GROMACS (*48*) and the CHARMM36m force field (*49*) with periodic boundary conditions.

Using the fibril ensemble as a reference, epitopes were predicted based on the consensus of three metrics that quantified local disorder, including loss of native contacts (ΔQ), increased root mean squared fluctuations (ΔRMSF), and increased solvent accessible surface area (ΔSASA). For a more detailed description of this approach for predicting epitopes, see ref.s (*28 –34*).

### Epitope scaffolding using cyclic peptides

As described in Results section: Prediction of tau epitopes, 3 regions emerge from the collective coordinates prediction as candidate misfolding-specific epitopes: ^315^LSKVT^319^, ^343^KLDFK^347^, and ^365^GGGN^368^ (see Fig. 6 A). The ^365^GGGN^368^ epitope was not pursued further due to it’s lack of sequence complexity and expected lack of immunogenicity. To scaffold a single instance of the predicted ^315^LSKVT^319^ or ^343^KLDFK^347^ epitope, we computationally constructed “glycindel” cyclic peptides following the procedure described in Ref. (*28*). Briefly, each cyclic peptide scaffold consists of the epitope flanked on the Nand C-termini by *n* and *m* glycines respectively. We refer to these scaffolds as cyclo(*C*-*G_n_*-(EPITOPE)-*G_m_*) or (*n, m*)EPITOPE, where *C* is a cysteine residue for conjugating a scaffold to carriers such as Keyhole limpet hemocyanin (KLH) in order to increase immunogenicity. Cyclic peptide construction is implemented computationally by mutating the corresponding flanking residues to glycine or cysteine in the native fibril structure using SCWRL4 (*59*). The topology of the cyclic peptide is obtained by head-to-tail backbone linkage using a locally written Python script (*28*), which is then energy minimized using a steepest descent algorithm in GROMACS (*48*).

**Figure 6:**
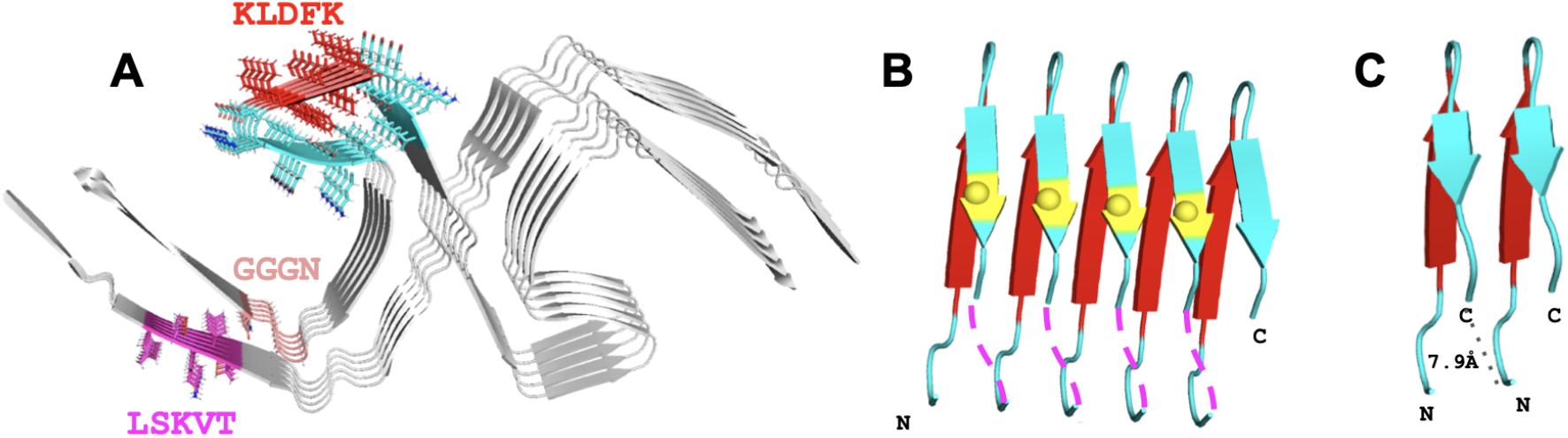
Modeling tau. *β***-helix structure.** (**A**) Tau fibril from Alzheimer’s disease brain (PDB ID: 5O3L) showing the side chains on the left protofilament in the figure for the predicted epitopes: ^315^LSKVT^319^ (magenta), ^343^KLDFK^347^ (red), and ^365^GGGN^368^ (light orange), and the tau339-354 fragment ^339^VKSEKLDFKDRVQSKI^354^ (non-epitope portion in cyan) that subsumes the ^343^KLDFK^347^ epitope. (**B**) Segments of five chains of the tau339- 354 fragment are connected together with interchain linkers, to form a *β*-helix structure. Magenta dash lines indicate how glycine tripeptide linkers connect the C-terminus of one chain to the N-terminus of the next chain, and S352 is shown in yellow spheres. (**C**) The 7.9Å distance between the C-terminus of one chain and the N-terminus of the next chain in the tau fibril (black dotted line). All images were rendered with PyMOL molecular visualization system (https://pymol.org)

### *β*-helix structure modeling

The initial *β*-helix structure was modeled using a repeating unit of 16 amino acid sequence spanning the tau fragment ^339^VKSEKLDFKDRVQSKI^354^ (tau339-354) (*43 –44*), which subsumes the ^343^KLDFK^347^ predicted epitope (see Fig. 6 A). Five tau339-354 repeating sequences were connected serially with glycine tripeptide (GGG) linkers to form a single chain, *β*-helix peptide with a total of 92 residues (Fig. 6 B).

The distance between the C*_α_* carbon of the C-terminal residue of tau339-354 on one chain and the C*_α_* carbon of the N-terminal residue of tau339-354 on the next consecutive chain in the tau fibril structure (PDB ID: 5O3L) was calculated to be ≈ 7.9Å (Fig. 6 C). We thus hypothesized that an ideal linker length would be about 3 amino acids, and investigated the distribution of distances in a non-redundant database of 26,900 structures (*60*). Figure 7 shows the distribution of distances between C*_α_* atoms of amino acids separated by (*i, i* + 3). We find several expected features, including peaks corresponding to *α*-helical and *β*-sheet secondary structure, along with an additional peak at ≈ 9Å separation corresponding to transitions from (*α, β*) secondary structures to loops or turns (*61*). These sequences were often initiated or terminated by glycines or prolines. The 7.9Å distance of our linkers (magenta bar labelled *ℓ* in Fig. 7) is in the middle of this distribution, supporting the hypothesis that a 3-amino linker was compatible in length and would not frustrate our designed structure.

**Figure 7:**
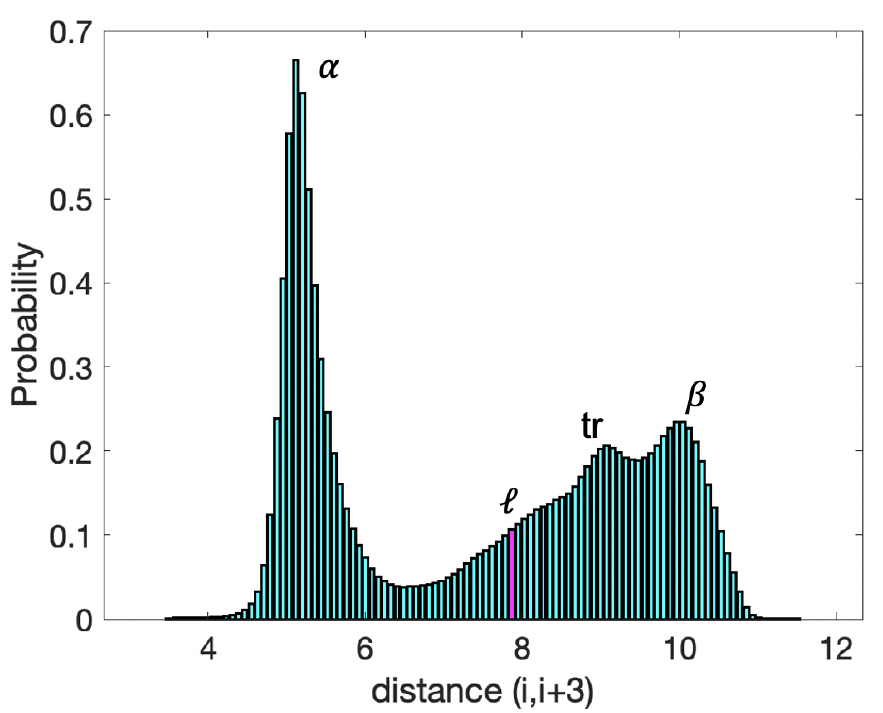
Distribution of. (*i, i* + 3) **distances in a non-redundant database of Protein Structures.** The 7.9Å distance between consecutive chains to be linked in our designed structure (*ℓ* in the plot) is well-within the distribution of distances between C*_α_* atoms separated by three amino acids (*i, i*+3). The distribution shows several features that are labelled in the figure: *α* (*α*-helical secondary structure), *β* (*β*-helical secondary structure), *tr* (transitional elements between secondary structure and loop or turn).

In order to further stabilize the designed scaffold, the serine at amino acid position 352 was mutated to cysteine for all but the last tau339-354 repeat, amounting to four S352C mutations in the structure (Fig. 1B). The S352C substitution introduces two disulfide bonds in the scaffold, which are introduced to improve the structural stability of the *β*-helix structure. The structure of the designed sequence was predicted using AlphaFold (*45*) (see Methods: Structure prediction with AlphaFold for further details), a deep learning algorithm for accurate protein structure prediction. Five models were generated by AlphaFold, and the model with the highest average confidence score that also formed the designed disulfide bonds between residues 14 and 33 and between residues 52 and 71 was selected as the initial model for the *β*-helix structure.

To further improve stability of the initial model, the structure of the C-terminal repeat sequence was remodelled to form an *α*-helix structure using the RosettaRemodel algorithm (*62*), while imposing the constraint that the distance between the Nand the C- termini C*_α_* atoms of the whole *β*-helix structure is ≈ 3.5Å (see Fig. 3 D). This remodeling step creates an interaction interface between the C-terminal *α*-helix and the GGG linkers. Furthermore, the remodeling allows for head-to-tail cyclization of the scaffold if needed, since the termini are now in spatial proximity.

### *β*-helix sequence design and structure optimization

Starting from the designed *β*-helix structure with C-terminal *α*-helix discussed above, the Rosetta protein design software (*46 –47*) was used to further optimize the sequence and overall structure of the modeled tau *β*-helix. PyRosetta (*63*) (Linux release 311 64Bit) was used for all Rosetta designs. The FastDesign protocol was used to pack all residues, and to apply sequence design to (only) the four linkers and the C-terminal *α*-helix residues, using the REF15 scoring function (*64*). All 20 naturally occurring amino acid residues were allowed in the designed residue positions. A MoveMapFactory was set up to allow the minimization of all bond lengths, bond angles, and torsion angles. The design protocol was repeated ten thousand times to generate 10,000 decoys of the *β*-helix structure. The Python code for structure remodeling, sequence design, and structure optimization using Rosetta can be found on github (see Methods: Code availability).

### Structure prediction with AlphaFold

DeepMind’s AlphaFold (version 2), a deep learning structure prediction algorithm that showed the best performance at CASP14 by a significant margin (*45 –65*) (https://github.com/deepmind/alphafold), was installed locally and trained, searching through the full database current as of January 01, 2020, and the model obtained was used to predict the structures of selected Rosetta designed *β*-helix scaffold decoys from their primary sequence alone.

### Mechanical unfolding simulations

To determine mechanical stability of the tau *β*-helix scaffold, unfolding simulations were performed. Ten independent forced unfolding simulations were performed along the reaction coordinate *Q* (the fraction of native contacts) over the course of 100 ns each, using PLUMED (*66*), OpenMM (*67*) simulation engine, and the CHARMM36m force field (*49*). A native contact was defined to be between heavy atoms within 0.45 nm, belonging to residues *α* and *β* with a sequence separation |*α* − *β*| *>* 3. The reaction coordinate *Q* of an arbitrary conformation *X* is defined as follows (*68*):

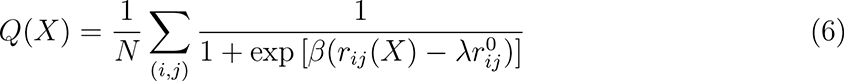

where the sum runs over N pairs of native contacts (*i, j*), *r_ij_*(*X*) is the distance between atoms *i* and *j* in conformation *X*, and *r*^0^ is the native distance between atoms *i* and *j*.

The parameters *β* and *λ* were taken to be 50 nm*^−^*^1^ and 1.5 respectively. A time-dependent harmonic potential *V* (*Q, t*) (Eq. 3) was applied to move the center of the bias linearly from *Q*_0_ = 1 to *Q*_0_ = 0 over the simulation time. The spring constant *k* in Eq. (3) was taken to be 10^7^ kJ·mol*^−^*^1^.

### Hamiltonian replica exchange molecular dynamics

Hamiltonian replica exchange molecular dynamics (HREMD) simulations (*69 –70*) were performed to determine the free energy surface (i.e. potential of mean force (PMF)) along the fraction of native contacts *Q* (Eq. 6). The set up consists of 100 replicas with time-independent harmonic restraints centered at positions equally spaced between *Q* = 1 and *Q* = 0. The strength of the harmonic restraint *k* was taken to be 10^5^ kJ·mol*^−^*^1^ for all replicas for the production run (80 ns for each replica). This value of harmonic restraint corresponds to an RMS values of Δ*Q* = 0.009 (this was about twice the 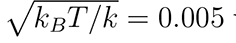 value expected from equipartition theorem). Before the production run, the protein must be unfolded and equilibrated in each harmonic potential restraint. The unfolding run is performed in 10 ns for each replica, and the system in each of the 100 harmonic potentials is equilibrated for 160 ns in the equilibration run.

The initial configurations before unfolding in each of the replicas were obtained by seeding all 100 replicas with the Rosetta-designed native structure, and applying the time-dependent harmonic restraints with corresponding center linearly decreased from *Q* = 1 to *Q* = *Q*_0_*_i_* during the unfolding run. Here, *Q*_0_*_i_*, 0 *< i <* 99, is the center of the harmonic restraint used for each replica during the equilibration and production runs. *Q*_0_*_i_* is different for each of the replicas, increasing linearly from 0.01 to 1 see Fig. 8). Because the interactions stabilizing the protein are stronger at higher nativeness, in practice we linearly decrease the strength of the harmonic restraints during the unfolding run as a function of time, from a spring constant of *k* = 10^7^ to *k* = 10^5^ kJ·mol*^−^*^1^. The spring constant then remains fixed at *k* = 10^5^ kJ·mol*^−^*^1^ during the equilibration and production runs. Exchanges between neighbouring replicas were attempted every 1 ps (500 steps). The simulation, including the initial and the production runs, ran for 250 ns per replica for a total of 25*µ*s for all replicas. Conformations were sampled at 10 ps intervals from each replica in the production run and used to determine the unfolding free energy profile along *Q* by solving the Multistate Bennett Acceptance Ratio (MBAR) equations (*71 –72*).

**Figure 8:**
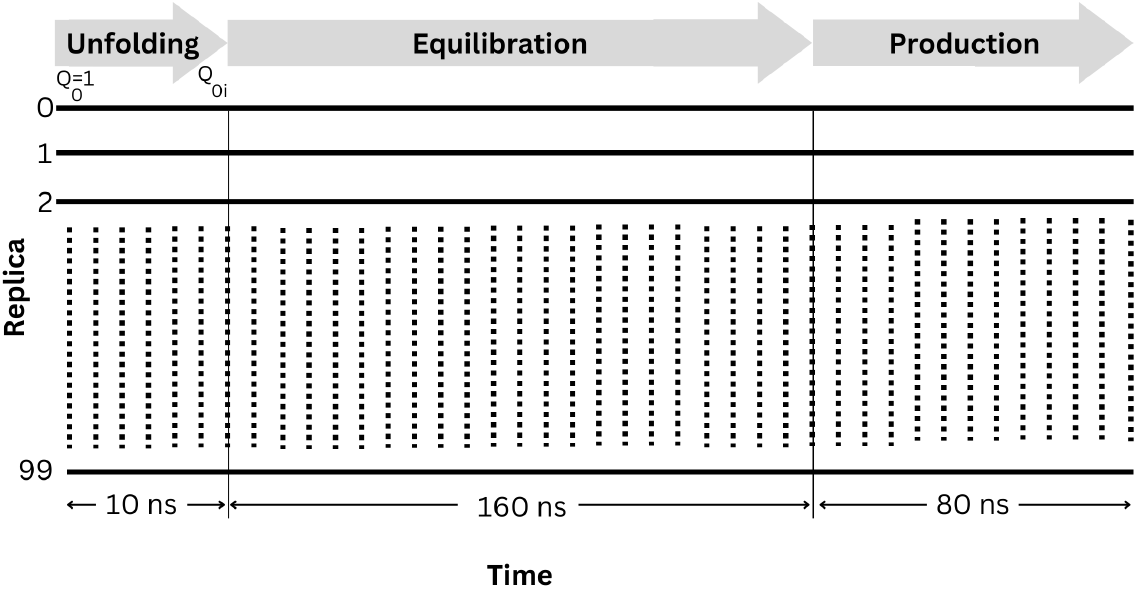
Setup used for hamiltonian replica exchange molecular dynamics (HREMD) simulations. The HREMD set up consists of 100 replicas (horizontal lines), each simulated in an unfolding run to different degrees of unfolding *Q*_0_*_i_* for 10 ns, followed by an equilibration run for 160 ns with each replica in its own harmonic potential centered at *Q*_0_*_i_*, followed by a production run in the same harmonic potential for 80 ns.

### Sampling conformational ensembles

All-atom molecular dynamics (MD) simulations were performed in explicit solvent (TIP3P water model, 150 mM NaCl) at a temperature of 300K and pressure of 1 bar using GROMACS (*48*) and CHARMM36m force field (*49*) to generate ensembles for tau fibril, tau oligomer, tau monomer, cyclic peptide scaffolds, and *β*-helix scaffold.

#### Fibril ensemble

The fibril ensemble was generated by performing 30.1 ns MD simulation starting from a tau fibril structure (PDB ID: 5O3L). Conformations were sampled at equal time intervals of 1 ns for a total of 30.1 × 10 × 10 = 3010 single-chain conformations (there are 10 chains in the system) in the fibril ensemble.

#### Oligomer ensemble

The oligomer was modeled as a partially unfolded fibril. Simulations were performed as described in the Prediction of epitopes section. Conformations were sampled at equal time intervals of 1 ns from the last ≈ 42 ns of ten independent biased simulations of 10 chains, for a total of 42 × 10 × 10 = 4200 single-chain conformations in the oligomer ensemble *Monomer ensemble:* The free energy landscape of an intrinsically disordered protein (IDP) such as tau is rugged and weakly funneled. Therefore, it is very challenging to generate an equilibrium ensemble for an IDP through direct MD simulation. The strategy that we have used previously in ref. (*28 –73*) was therefore employed. In brief, two steps are involved. Firstly, a highly diverse conformational ensemble with 10,000 configurations was generated by using the pivot algorithm (*74 –75*), along with crankshaft moves (*76*) that randomize torsion angles between two randomly-selected fixed backbone atoms. Secondly, each feasible conformation generated was energy minimized, and then a 3 ns MD simulation was performed. The last conformation from each simulation was collected to obtain a total of 7166 conformations in the monomer ensemble.

#### Cyclic peptide ensembles

Each cyclic peptide scaffold was simulated for 300 ns using conventional MD. Conformations were then sampled at equal time intervals of 100 ps from the last 250 ns of the simulation, for a total of 2500 conformations in each cyclic peptide scaffold ensemble.

#### β-helix scaffold ensembles

For the *β*-helix peptide ensembles, we performed 100-replica HREMD simulations for a total of 25 *µs* (see Methods: Hamiltonian replica exchange molecular dynamics). A number N*_i_* of conformations determined from Eq. 7 were sampled randomly from the *i*th replica, so that replicas were Boltzmann-weighted:

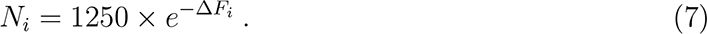

Here, Δ*F_i_*is the relative free energy (in k*_B_*T) of the *i*th replica (see Fig. 4 D). A total of 2393 conformations were sampled for each of four instances of KLDFK motif in the *β*-helix scaffold.

### Comparing conformational ensembles

Conformational ensembles are compared using Jensen-Shannon Divergence (JSD) (*77 –78*) and Embedding Depth (D) (*28*) measures. JSD was implemented using the ENCORE software (*79*). JSD is a symmetrized version of the Kullback–Leibler divergence (*80*), *D_KL_*, which is a measure of the difference between two probability distributions *P* and *Q*, defined by

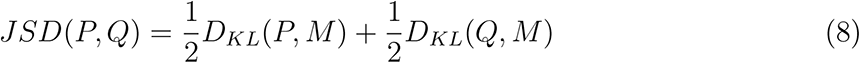

where

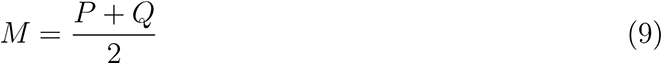

and

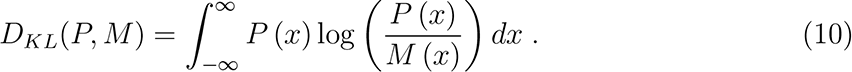

The probability distribution representing an ensemble is obtained by creating a root mean squared deviation (RMSD) matrix of all structures in all ensembles to be compared. The dimension of this high-dimensional RMSD matrix is then reduced using the Stochastic Proximity Embedding (SPE) method (*81*). Since the value of the JSD depends on the dimension of the SPE-reduced space, weighted average JSD values from 3D to 11D are used, with weights taken as the inverse of the SPE residuals in each dimension (this weights higher dimensions more strongly). The maximum value of JSD in Eq. 8 is log 2. JSD was then normalized to lie between 0 (for identical ensembles) to 1 (for entirely different ensembles) by dividing by log 2.

The Embedding Depth 𝒟*_Q|P_* or D_Q-in-P_, when used to compare two conformational ensembles represented by probability distributions *P* and *Q*, quantifies the extent to which the ensemble *Q* is subsumed by ensemble *P* . For a point **x** = **x_o_** in a distribution *P* (**x**), the embedding depth 𝒟*_δ_*_(_**_x_***_−_***_xo_**_)_*_|P_* (Eq. 11) is defined as the fraction of the distribution that has a lower probability than point **x** = **x_o_** (*28*):

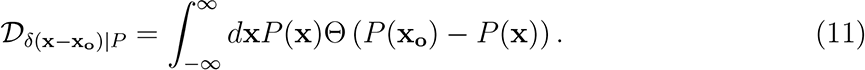

In Eq. 11, Θ(*P*) is the Heaviside step function, which returns 1 if *P >* 0, otherwise 0.

The Embedding Depth of one distribution *Q*(**x**) within another *P* (**x**) can be found by integrating Eq. 11 over the distribution *Q*(**x**):

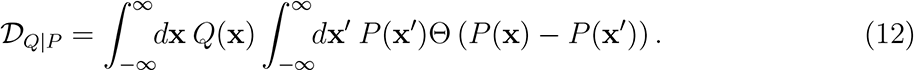

The probability distribution representing an ensemble for Embedding Depth calculation is also obtained by creating a RMSD matrix of all structures in all ensembles to be compared. The dimension of this high-dimensional RMSD matrix is however reduced using the Multidimensional Scaling (MDS) method (*82 –84*). Since the value of 𝒟 also depends on the dimension of the MDS-reduced space, a weighted average of 𝒟 values from 3D to 11D are used, with weights taken as the inverse of the MDS residuals in each dimension (similar to JSD above, this applies more weight to higher dimensions). The Embedding Depth between two distributions is in general non-reciprocal, in that 𝒟*_Q|P_* ≠ 𝒟*_P |Q_*. The mode of a distribution has the maximum value of 𝒟 = 1 while two entirely different ensembles have the minimum value of 𝒟 = 0. The embedding depth of a distribution within itself is 𝒟*_P |P_* = 1*/*2 (*28*).

### Epitope robustness analysis

The SASA (averaged across chains) is plotted as a function of residue position, for each of the five tau fibril structures in Fig. S7. A rolling average window of 5 amino acids was applied. The pairwise local distance test (lddt) (*38*) was used to compare PDB fibril structures to determine their similarity (Fig. S7 G).

### Protein cleavage analysis

Cleavage analysis was performed by the Procleave server (*55*), with default parameters on all proteases. The proteolytic propensity of the designed *β*-helix scaffold structure was compared to ubiquitin (PDB 1UBQ (*85*)) as a negative control. Plots were made of the top 10 cleavage site scores for 27 specific proteases (Fig. S8).

### Aggregation propensity analysis

Aggregation propensity analysis was performed using the AGGRESCAN3D server(*56*) with default parameters. The designed *β*-helix scaffold structure was compared to ubiquitin (PDB 1UBQ (*85*)) as a negative control, and transthyretin (PDB 1TFP (*86*)) as a positive control for an known aggregation-prone protein (*57*). Plots of the aggregation score vs. residue index, along with the mean aggregation score and the fraction of residues that have an aggregation score larger than zero, are shown in Fig. S9.

### Immunogenicity analysis

Predicted immunogenicity score was obtained from the Epitopia server. (*39*) Fig. S10 plots the immunigenicity for each residue on a scale from 1 to 5, for the tau fibril structure (PDB 5O3L). The number plotted in Fig. S10 is averaged over the 10 chains in the fibril structure.

### Design method for the cyclic CPPPPKLDFKGPGG scaffold

The centroid structures of the biased fibril ensemble, as well as the monomer ensemble of the KLDFK segment, were both identified. Both centroid structures were scaffolded using a (4,4) glycindel scaffold, consisting of four glycines on both the C- and N-terminus. To identify Gly→Pro mutations that would reinforce conformational similarity to the centroid structure of the biased fibril, while avoiding similarity to the centroid structure of the monomer ensemble, the CRISPro software (*87*) was employed. This analysis led to the mutations at five specific proline mutation sites, modifying the sequence from CGGGGKLDFKGGGG to CPPPPKLDFKGPGG. A cyclic peptide with this latter sequence was used in immunizations to obtain the monoclonal antibodies described in Methods: Surface plasmon resonance (SPR) experiments.

### Peptide Synthesis

Peptide synthesis was performed by LifeTein, LLC. (Hillsborough, NJ, USA) following standard manufacturing procedures. Mass spectrometry QC indicated the correct molecular weight to within a protonation state, and was consistent with disulfide bond formation (Theoretical value 11336.99, Observed on MS at 11336.05). HPLC indicated a purity value at 85.04% by peak area. Designed constructs were conjugated to keyhole limpet hemocyanin (KLH).

### Surface plasmon resonance (SPR) experiments

Several monoclonal antibodies have been obtained previously from mice immunized with the Tau cyclic peptide sequence CPPPPKLDFKGPGG. These include antibodies 4E4, 3E2, 10B10, 6H3, 7H6, 10D4, 7E5, and 2C6 (see Figure 5). An isotype control (mIgG1) was also used for comparison.

Stable Tau Oligomers (consisting primarily of trimers and pentamers) were generated from recombinant full-length human tau monomers (wild-type tau—441 amino acids, 2N4R, *>*97% purity), were obtained from SynAging SAS (Vandœuvre-lès-Nancy, France).

Tau *β*-helix construct reduction was performed by incubating 0.8mg/ml (90*µ*M) sample with 10mM DTT (90X molar excess), and 2M EDTA for 30 minutes at room temperature. Approximately 400 RUs of oxidized and reduced *β*-helix construct were immobilized for SPR binding studies. Antibodies were diluted to 150*µ*g/ml and injected over immobilized *β*-helix constructs or oligomers.

### Code availability

A Python script for remodelling and design of the tau beta-helix protein scaffold is available at https://github.com/PlotkinLab/Beta-Helix-Design

## Conclusion

In this paper, we have presented a novel approach for scaffolding epitopes in order to target toxic forms of proteins implicated in protein misfolding diseases. This approach entails the design of scaffolds that connect several chains in an aggregated structure by designed linkers, to incorporate multiple repeats of a target epitope. This construction is intended to guide the conformational ensemble explored by the epitope towards those conformations present in aggregated conformations, which we hypothesize would present the epitope in disease-associated forms of the underlying protein. The prediction of what epitopes to scaffold in this manner would depend on the application; in our context, the epitopes were predicted to be those on the accessible surface of an oligomer.

The method was applied to tau, a microtuble-binding protein whose loss of function and tangle formation in neurons is associated with many neurodegenerative diseases, including Alzheimer’s disease and other tauopathies. First, oligomer-selective tau epitopes were predicted by stressing the fibril to induce a partially disordered fibril, which serves as a predictive model for epitopes that are likely to be exposed on the surface of oligomers.

Four copies of the predicted tau ^343^KLDFK^347^ epitope were incorporated in the final design of a *β*-helix scaffold. The structure of the designed scaffold was validated with AlphaFold, and it is gratifying to see that the Rosetta-designed structure and AlphaFold-predicted structure are nearly identical, with a C*_α_* RMS deviation of 1.39 Å. *In silico* characterization of the designed scaffold shows that it is predicted to be thermodynamically stable, which is an important property of a molecule intended for use as an immunogen.

Furthermore, we showed that for this epitope in tau protein, the multi-epitope scaffolding approach is predicted to be better in discriminating models of abnormal forms of tau from isolated tau monomer, than single-epitope scaffolding approaches, such as in cyclic peptide “glycindel” scaffolds. The multi-epitope scaffolding approach introduced here can straightforwardly be applied to proteins implicated in other protein aggregation-related diseases, such as tau protein in chronic traumatic encephalopathy (CTE), *α*-synuclein in Parkinson’s disease, SOD1 in amyotrophic lateral sclerosis (ALS), TDP43 in frontotemporal dementia (FTD), and Huntingtin in Huntington’s disease.

The experimental characterization included here showed that immunological antibodies raised to a cyclic peptide of the KLDFK epitope could bind to the *β*-helical construct, that their binding is weakened by loss of structure of this construct, and interestingly, that their strength of binding to the *β*-helical construct is correlated with their strength of binding to *in vitro* oligomers of tau. This supports the notion that the *β*-helix construct may present an oligomer-selective conformational epitope, and may thus be useful as a potential immunogen. It is currently unknown how effective such an immunogen would be in the human immune system to raise an oligomer-selective antibody response.

## Supporting Information Available

Supplementary Information is available at the of the manuscript.

1.(PDF) Figures S1—Snapshots of tau fibrils; S2—“Fireplots” of the change in SASA upon stressing the fibril; S3—“Fireplots” of the loss of native contacts upon stressing the fibril; S4—“Fireplots” of the change in RMSF upon stressing the fibril; S5—Rosetta energy and AlphaFold confidence for designed *β*-helix structures; S6—Instability of 4-repeat *β*-helix scaffold; S7—Commonly high SASA of KLDFK motif among tau fibrils; S8—Protease analysis of *β*-helix scaffold; S9—Aggregation propensity of *β*helix scaffold; S10—Immunogenicity propensity of *β*-helix scaffold; S11—Secondary structure evolution of *β*-helix scaffold; Table S1—Models of tau *β*-helix scaffold
2.A github repository link to the code for *β*-helix scaffold design is at https://github.com/PlotkinLab/Beta-Helix-Design

## Author information

### Corresponding author

Steven S. Plotkin — Department of Physics and Astronomy, The University of British Columbia, Vancouver, BC, V6T 1Z1, Canada; Genome Science and Technology Program, The University of British Columbia, Vancouver, BC, V6T 1Z1, Canada.

### Authors

Adekunle Aina — Department of Physics and Astronomy, The University of British Columbia, Vancouver, BC, V6T 1Z1, Canada.

Shawn C.C. Hsueh — Department of Physics and Astronomy, The University of British Columbia, Vancouver, BC, V6T 1Z1, Canada.

Ebrima Gibbs — Djavad Mowafaghian Centre for Brain Health, University of British Columbia, Vancouver, BC V6T 1Z1, Canada.

Xubiao Peng — Department of Physics and Astronomy, The University of British Columbia, Vancouver, BC, V6T 1Z1, Canada; Center for Quantum Technology Research, School of Physics, Beijing Institute of Technology, Beijing, China 100081.

Neil R. Cashman — Djavad Mowafaghian Centre for Brain Health, University of British Columbia, Vancouver, BC V6T 1Z1, Canada; ProMIS Neurosciences, Cambridge, MA 02142, USA

### Author Contributions

*Conceptualization:* A.A. and S.S.P.; *Methodology:* A.A. and S.S.P.; *Coding—β-helix design:* A.A.; *Software:* A.A., S.C.C.H., X.P., and S.S.P.; *Data analysis:* A.A., S.C.C.H., X.P., and S.S.P.; *Visualization:* A.A., S.C.C.H., X.P., and S.S.P.; *Experiment—design and method:* E.G., N.R.C., and S.S.P.; *Experiment—data analysis:* E.G., N.R.C., and S.S.P.; *Writing— original draft:* A.A.; *Writing—revision, review and editing:* S.S.P., A.A., and S.C.C.H.; *Supervision:* S.S.P. and N.R.C.; *Funding acquisition:* S.S.P., A.A., and N.R.C.

## Acknowledgement

We thank Santanu Sasidharan for his help in rendering the experimental data.

## SUPPLEMENTARY INFORMATION

**Figure S1:**
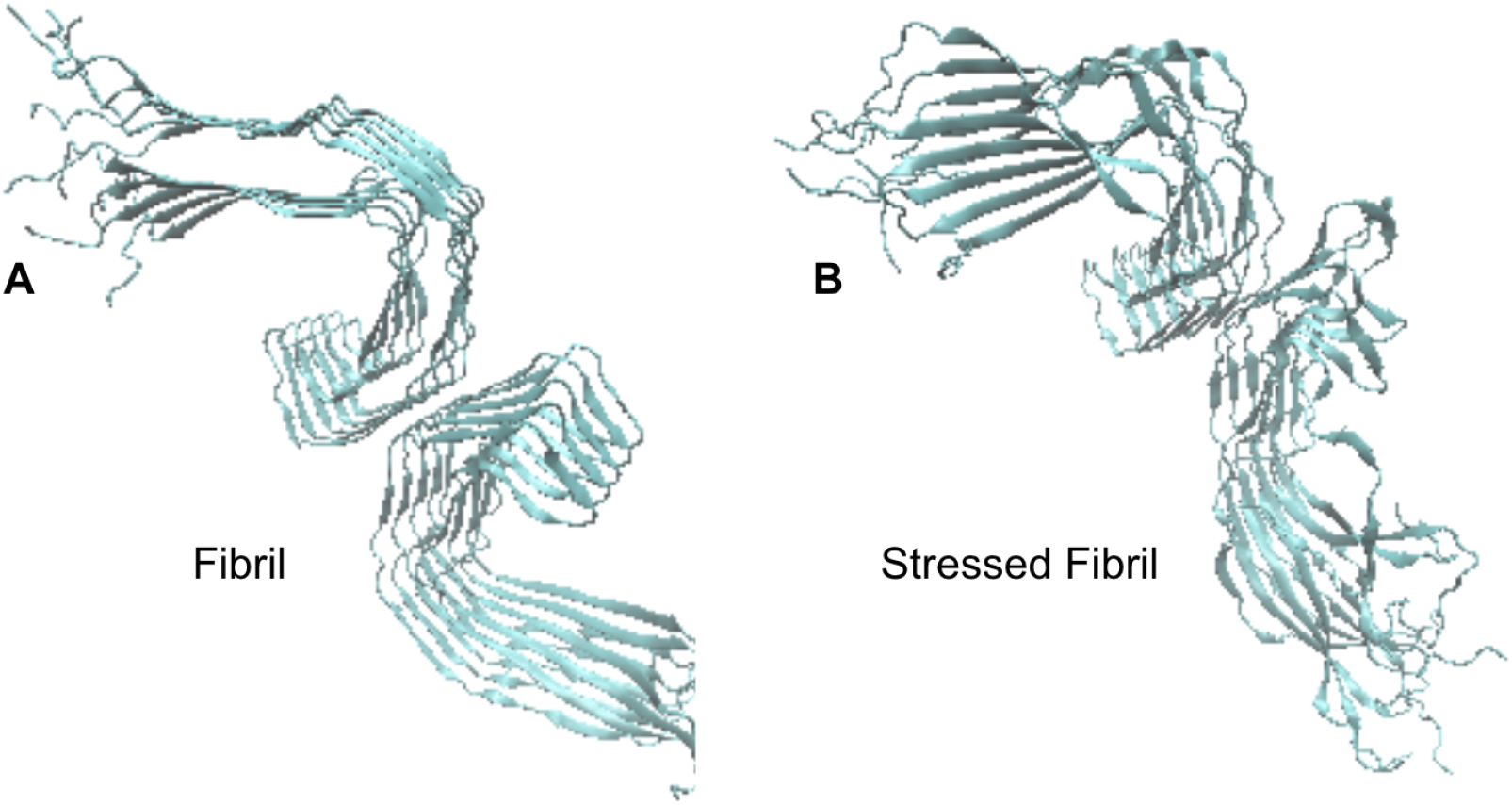
Snapshots of tau fibrils (PDB ID: 503L). (A) Tau fibril before introducing an unfolding biasing potential and **(B)** after biasing the fibril to 60% of its native contacts. Images were rendered using the VMD molecular visualization system^8^ (http://www.ks.

**Figure S2:**
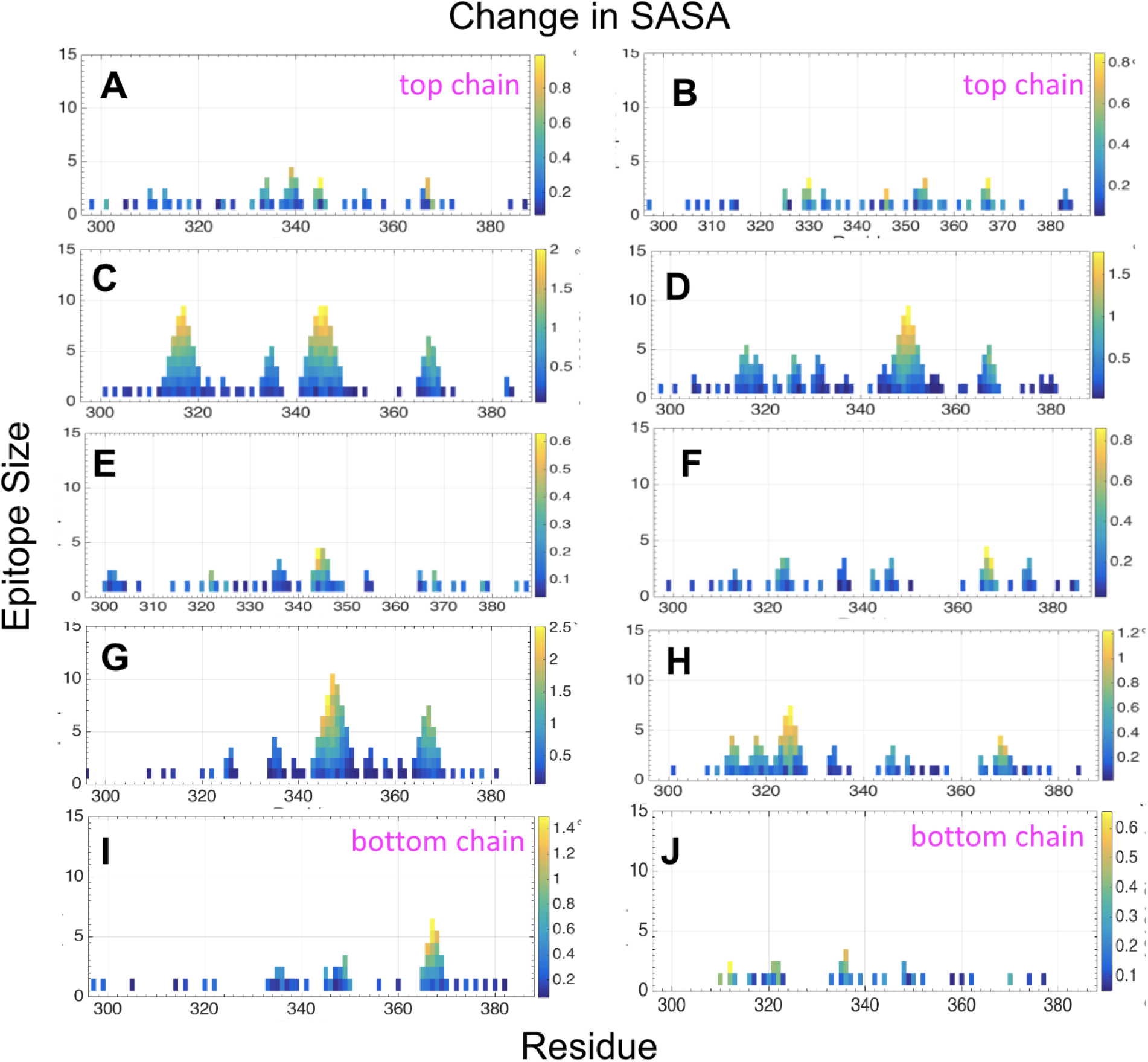
“Fireplots” of the change in solvent accessible surface area (SASA) upon stressing the fibril. The total SASA change after biasing the tau fibril (PDB ID: 503L) to 60% of its native contacts for **(A)** chain A, **(B)** chain B, **(C)** chain C, **(D)** chain D, **(E)** chain E, **(F)** chain F, **(G)** chain G, **(H)** chain H, **(I)** chain I, and **(J)** chain J. The *x*-axis indicates the central residue index of a predicted epitope. The central left residue is used for even residue-length epitopes. The *y*-axis indicates the number of residues in the epitope. So for example the largest predicted epitope in chain A is only 4 residues, left-centered on residue V339 and consisting of sequence _338_EVKS_341_, which is the highest point in panel A. The color coding gives the total ΔSASA, averaged over all simulations, of the group of underlying residues characterized by the size on the *y*-axis and center position on the *x*-axis. Only epitopes that satisfy ΔSASA *>* 0 for all residues in the epitope in at least 8 of 10 independent simulations were considered to be reliably exposed.

**Figure S3:**
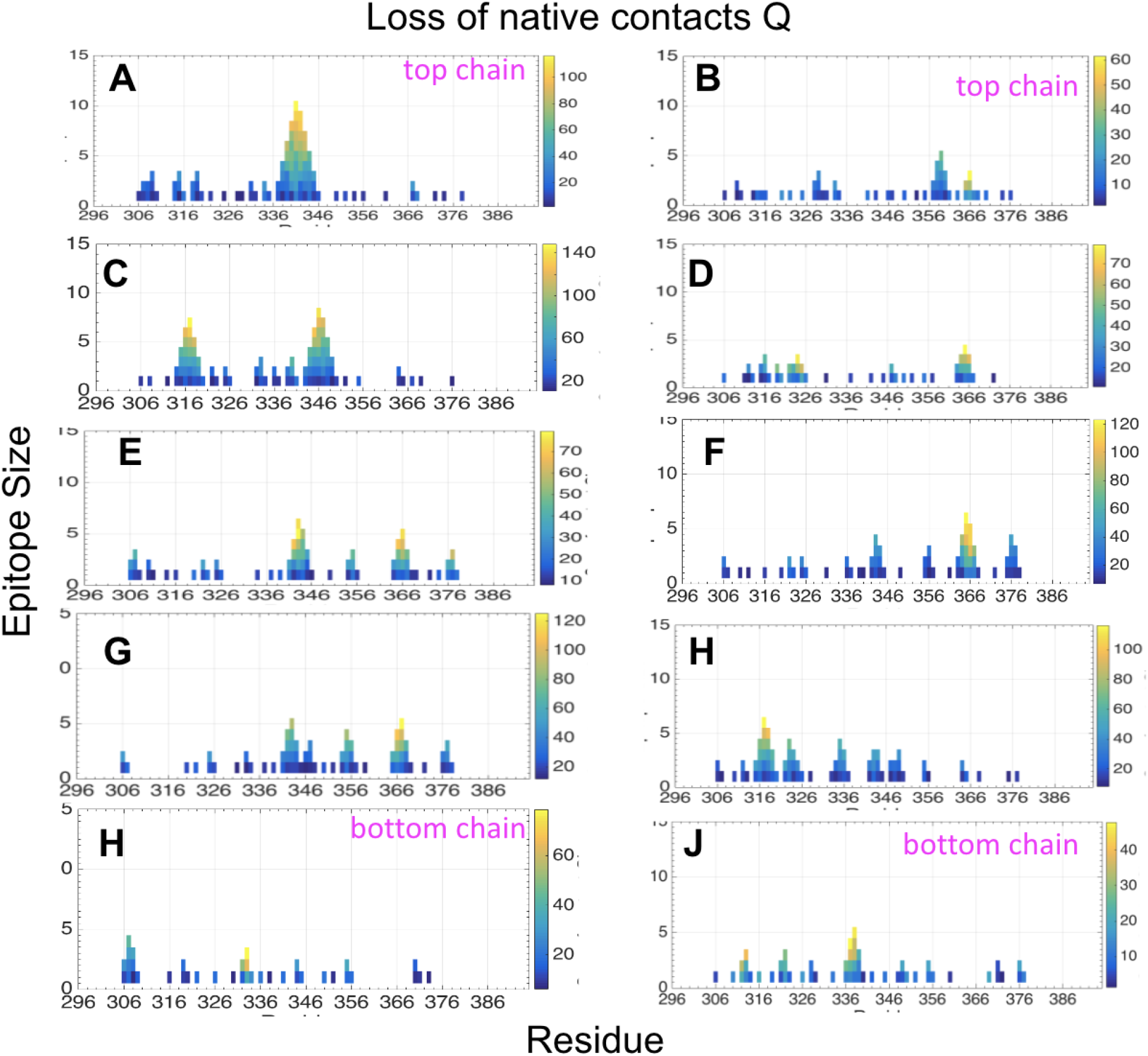
“Fireplots” of the loss of native contacts (Q) upon stressing the fibril. The loss of total contacts *Q* after biasing the tau fibril (PDB ID: 503L) to 60% of its “native” contacts for **(A)** chain A, **(B)** chain B, **(C)** chain C, **(D)** chain D, **(E)** chain E, **(F)** chain F, **(G)** chain G, **(H)** chain H, **(I)** chain I, and **(J)** chain J. The *x*-axis indicates the central residue index of a predicted epitope. The central left residue is used for even residue-length epitopes. The *y*-axis indicates the number of residues of the epitope. The color coding gives the total loss ΔQ averaged over the number of simulations of the group of underlying residues characterized by the size on the *y*-axis and center position on the *x*-axis. Only epitopes that satisfy ΔQ *>* 0 for all residues in the epitope in at least 8 of 10 independent simulations were considered.

**Figure S4:**
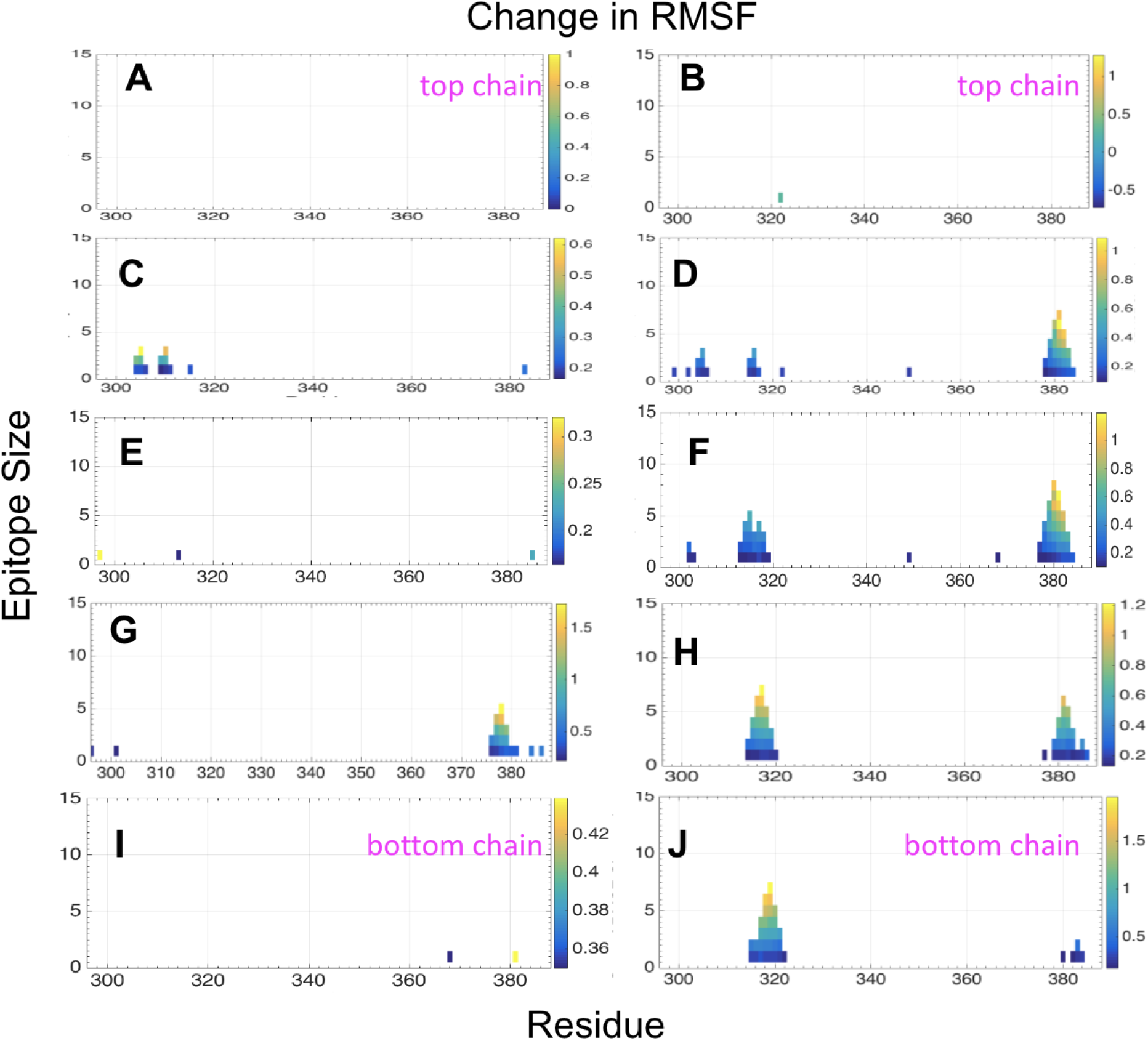
“Fireplots” of the change in root mean square fluctuations (RMSF), upon stressing the fibril. The total change in RMSF after biasing the tau fibril (PDB ID: 503L) to 60% of its native contacts for **(A)** chain A, **(B)** chain B, **(C)** chain C, **(D)** chain D, **(E)** chain E, **(F)** chain F, **(G)** chain G, **(H)** chain H, **(I)** chain I, and **(J)** chain J. The *x*-axis indicates the central residue index of a predicted epitope. The central left residue is used for even residue-length epitopes. The *y*-axis indicates the number of residues of the epitope. The color coding gives the total ΔRMSF averaged over the number of simulations for the group of underlying residues characterized by the size on the *y*-axis and center position on the *x*-axis. Because ΔRMSF is positive for nearly all residues after biasing, only epitopes that satisfy ΔRMSF *>* ⟨RMSF⟩ for all residues in the epitope in at least 8 of 10 independent simulations were considered.

**Figure S5:**
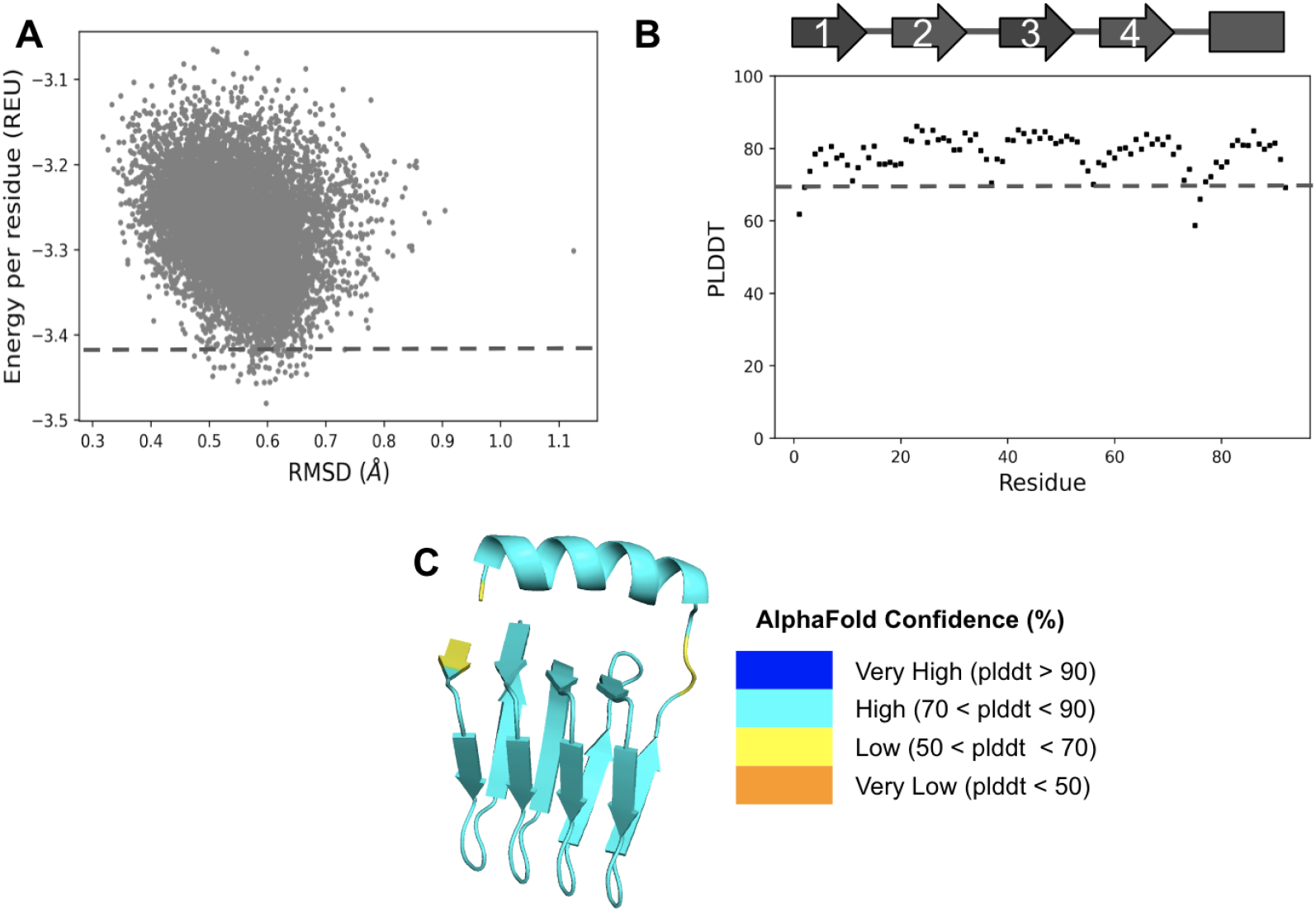
Rosetta energy and AlphaFold confidence for designed. *β***-helix structures. (A)** Rosetta^1,2^ energy per residue (in Rosetta energy units or REU) as a function of C*_α_* RMSD to the Rosetta remodeled structure, for 10, 000 designed decoys. Dashed line indicates low energy cuttoff of −3.41 REU, corresponding to 1% of decoys with energies below the cuttoff. **(B)** Per residue AlphaFold^3^ confidence (PLDDT) score for the top-ranked AlphaFold models as for the top ranked sequence. Dashed line indicates high confidence cuttoff of 70%. (Top) Secondary structure schematic for *β*-strands (arrow) representing tau339-354 (^339^VKSEKLDFKDRVQSKI^354^) repeats and remodeled *α*-helix (rectangle). The *β*-helix structure was rendered to show **(C)** 1st ranked per residue confidence scores and colour coded as very high confidence (blue), high confidence (cyan), low confidence (yellow), and very low confidence (orange). Image were rendered with PyMOL molecular visualization system (https://pymol.org)

**Figure S6:**
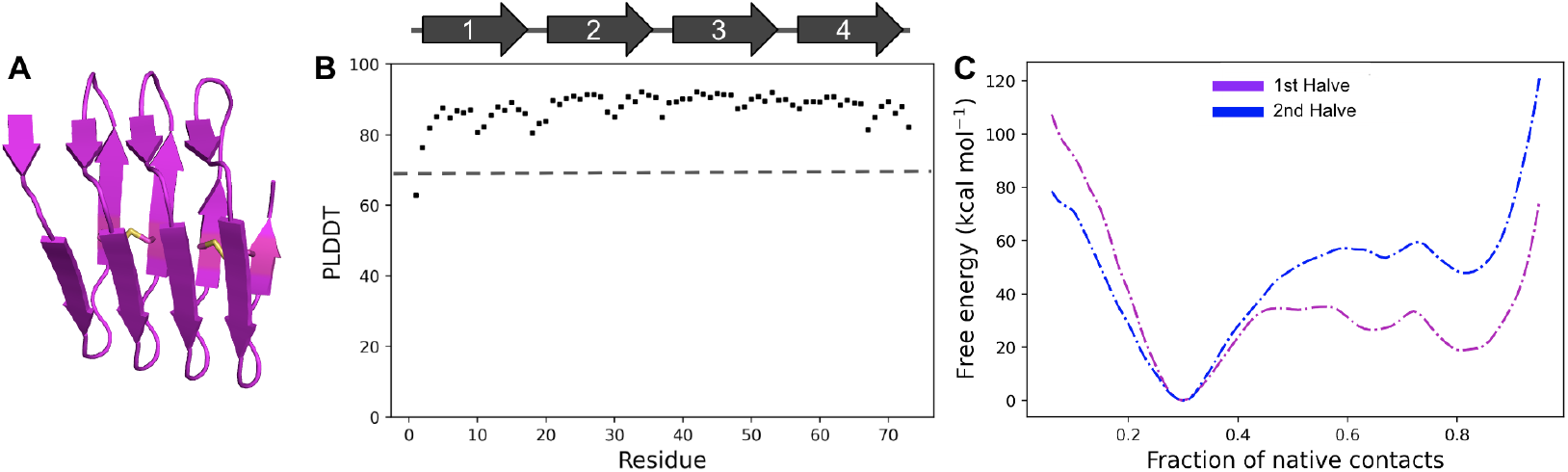
Instability of 4-repeat. *β***-helix scaffold. (A)** AlphaFold predicted structure and **(B)** Per residue AlphaFold confidence (PLDDT) score is high for the top-ranked AlphaFold model. **(C)** Unfolding free energy profile indicates that the global free energy minimum is the unfolded state. The first and 2nd halves of the simulation trajectories are shown. The Rosetta designed linkers for linker positions 1, 2 and 3 are GGG, AGN, and GPG, respectively. Image was rendered with PyMOL molecular visualization system

**Figure S7:**
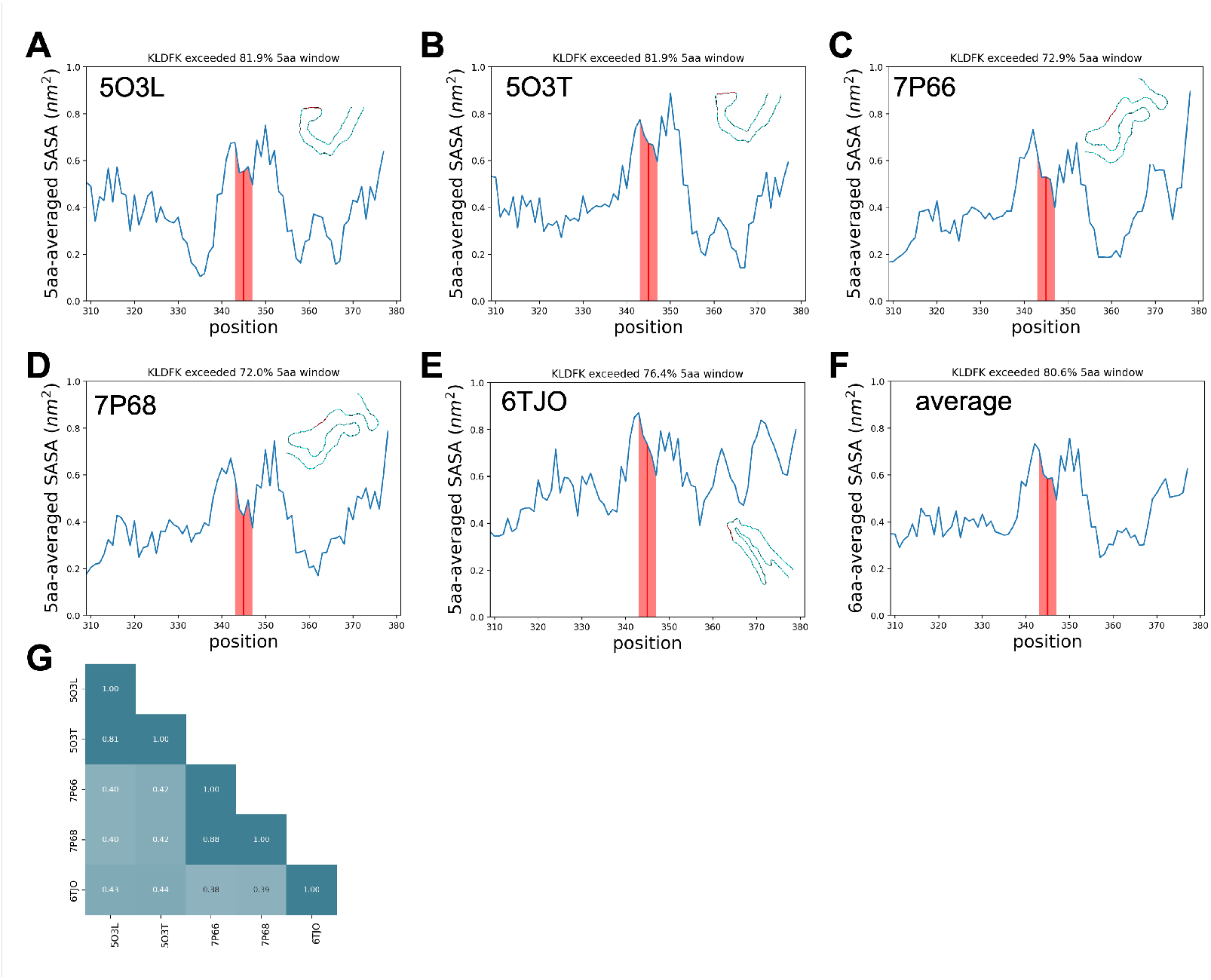
Commonly high SASA of KLDFK motif among tau fibrils. The collective coordinates-predicted tau protein epitope, KLDFK, is exposed to solvent in five tau distinct fibril structures: 5O3L, 5O3T, 7P66, 7P68, 6TJO. **(A—E)** Averaged SASA as a function of residue position. A rolling average window of 5 amino acids was applied. The window that contains KLDFK (residues 343-347), as indicated by the red line, exceeds more than 70% of the other windows in all fibril structures analyzed. The shaded region contains the rolling average values that partially contain residues 343-347. The epitope also tends to be flanked on either side by regions with even higher surface exposure. In each panel, a single chain of each fibril structure is aligned and rendered to show their structural polymorphism. **(F)** The average SASA across all 5 fibrils. The epitope region has high average SASA across the whole structured sequence, and appears poised to have higher surface exposure upon stress of the fibril. **(G)** The pairwise local distance test (lddt)^9^ between the fibril structures shows that 5O3L and 5O3T as well as 7P66 and 7P68 are mutually similar, resulting in 3 distinct structural classes.

**Figure S8:**
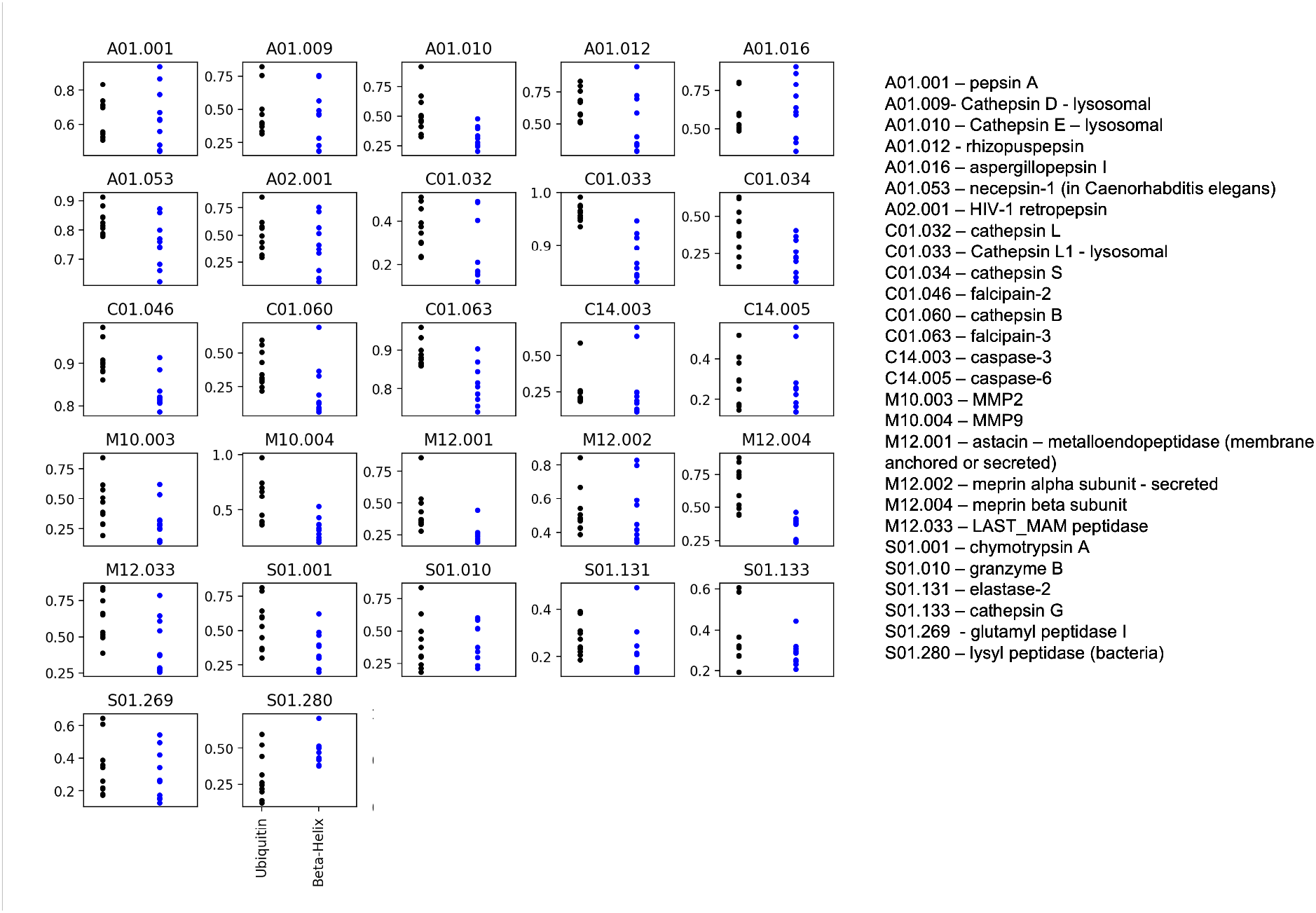
Protease analysis of. *β***-helix scaffold.** Protease cleavage analysis was performed using the designed *β*-helix scaffold structure (blue points) and the structure of ubiquitin (black points) (PDB 1UBQ^10^), which serves as a negative control. Each plot shows the top 10 cleavage site scores for a specific protease (as shown on the right). When comparing the highest scoring cleavage site between *β*-helix and ubiquitin for each protease, the *β*-helix sequence scored higher in only 8 out of the 27 proteases. Procleave^4^ server was used to perform cleavage analysis.

**Figure S9:**
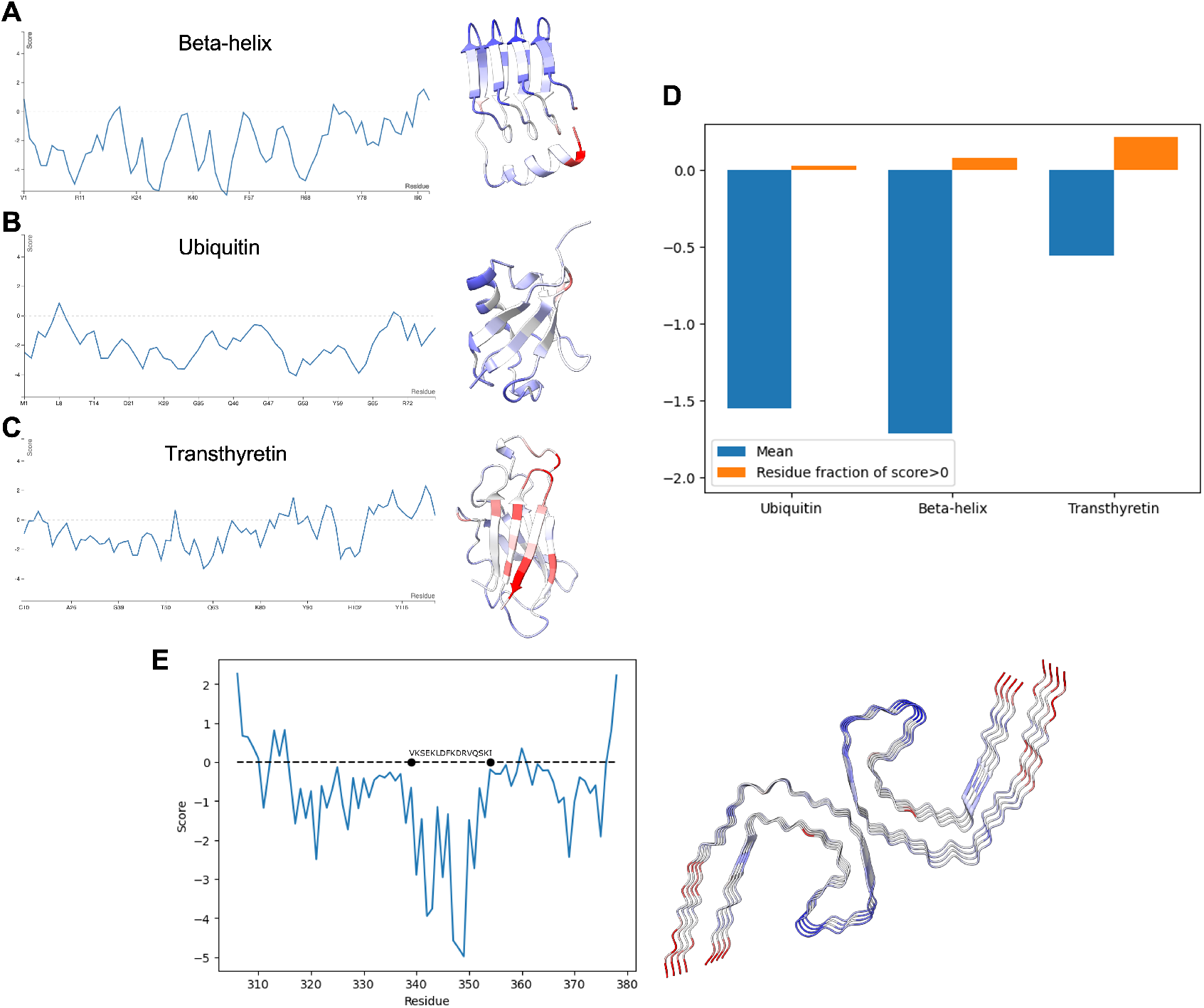
Aggregation propensity of. *β***-helix scaffold.** The AGGRESCAN3D^5^ server prediction for amyloid propensity *vs.* residue index for **(A)** *β*-helix scaffold **(B)** ubiquitin (PDB 1UBQ^10^), a negative control, and **(C)** transthyretin (PDB 1TFP^11^), a positive control. Residues are color-coded in the structures shown on the right by amyloid propensity. Panel **(D)** shows the mean of the aggregation scores, and the fraction of residues that have an aggregation score larger than 0, for the three proteins in panels A-C. **(E)** The aggregation score of the tau fibril (PDB 5O3L), as predicted by the AGGRESCAN3D server, has been shown by averaging the scores across eight chains A-H. The native region that was scaffolded to be *β*-helix (^339^VKSEKLDFKDRVQSKI^354^) is shown to be the least aggregation-prone region.

**Figure S10:**
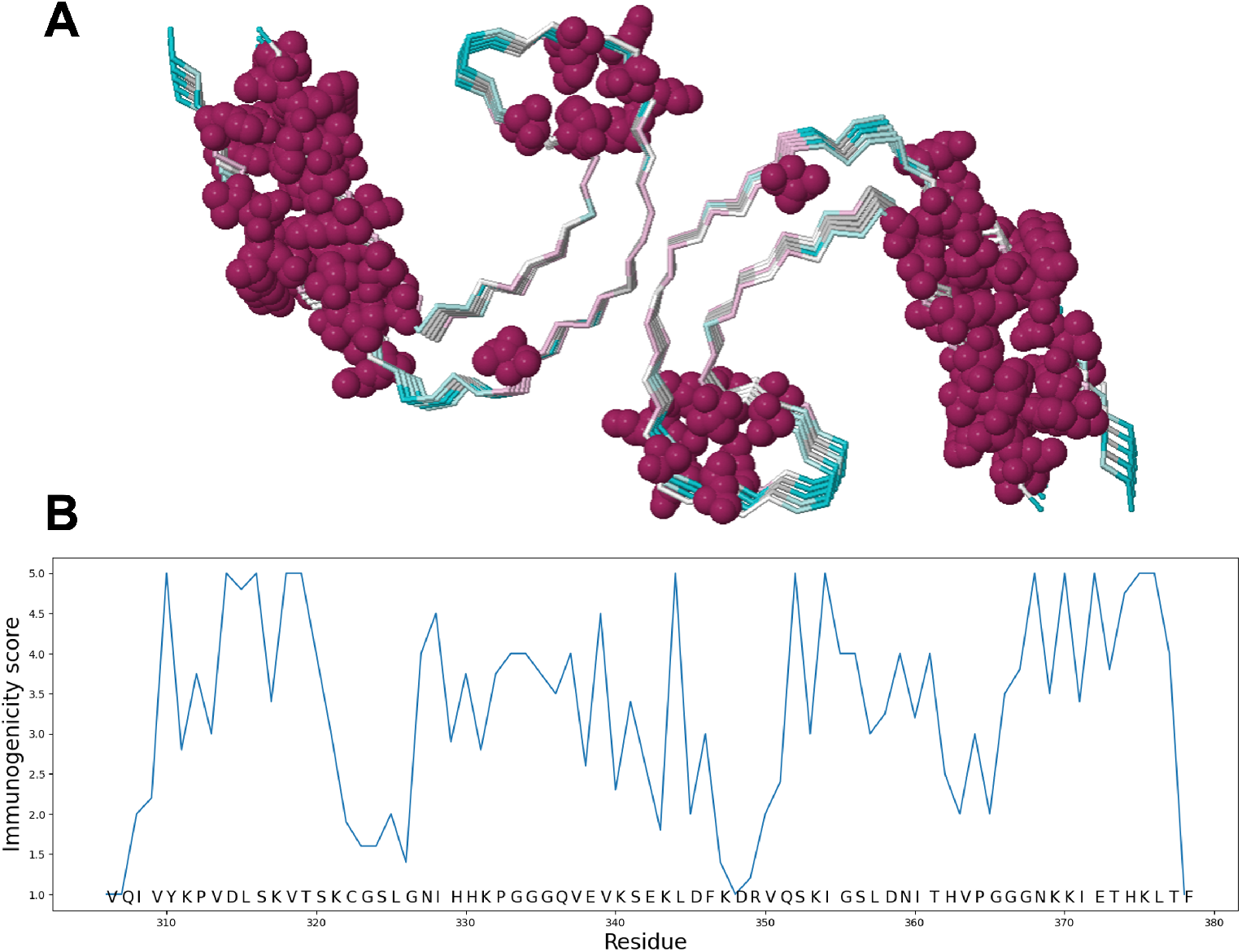
Immunogenicity propensity of. *β***-helix scaffold.** The Epitopia^6^ server was employed to predict an immunogenicity score ranging from 1 to 5 for each residue in Tau (PDB 5O3L). Panel **(A)** displays the sidechains of residues with an immunogenicity score of 5. Panel **(B)** demonstrates the immunogenicity scores averaged across each chain. ^343^KLDFK^347^ did not have a particularly high score. On the other hand, ^315^LSKVT^319^,, another misfolding-prone epitope predicted from the collective coordinates algorithm,^12^ did receive a high Epitopia score.

**Figure S11:**
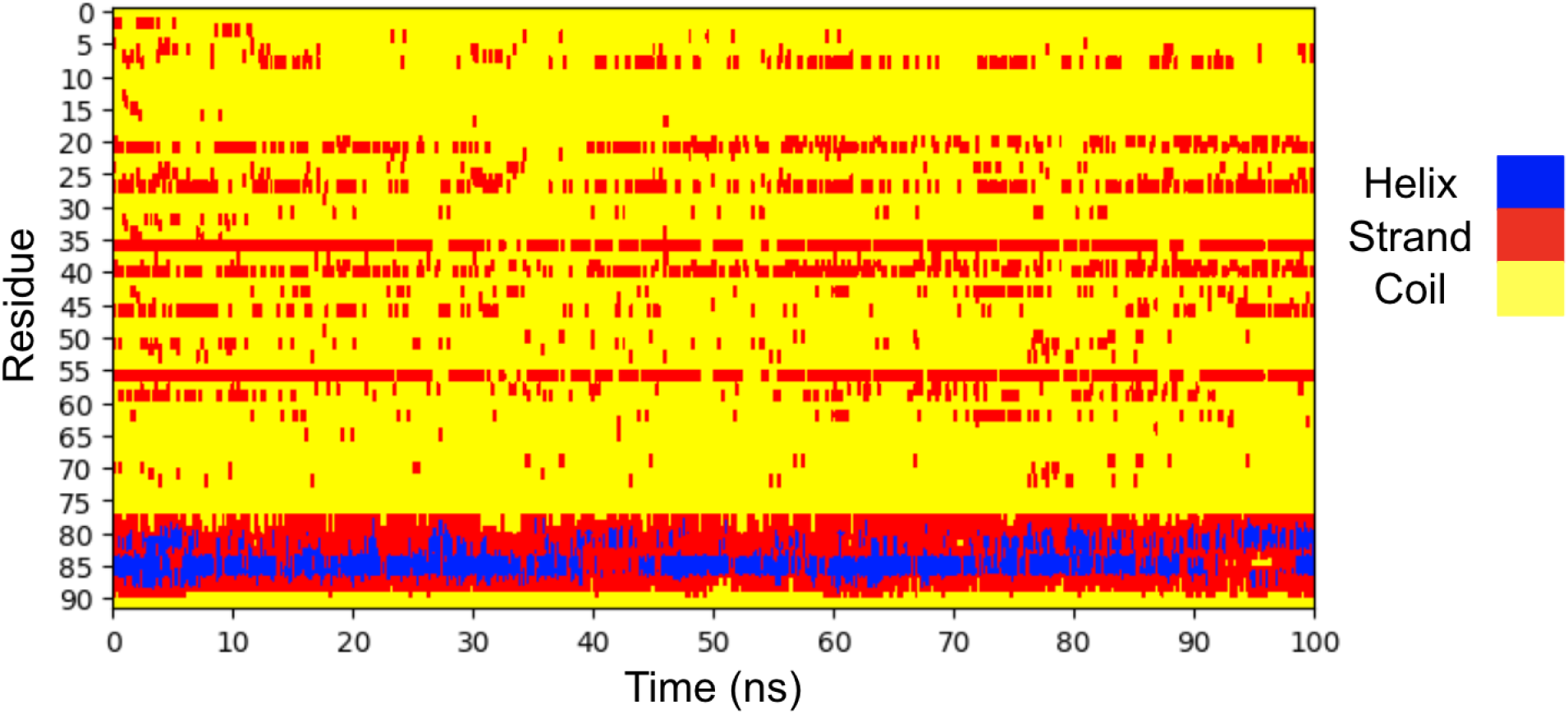
Secondary structure evolution of. *β***-helix scaffold.** The time evolution of the secondary structure of each amino acid residue in the designed *β*-helix scaffold over 100 ns molecular dynamics simulation. **(Blue)** Helix: either *α*-helix, 3/10-helix or *π*-helix, **(Red)** Strand: either extended *β*-strand or isolated *β*-bridge, and **(Yellow)** Coil: either loop, bend or hydrogen-bonded turn. Secondary structure was assigned by DSSP.^7^

**Table S1:**
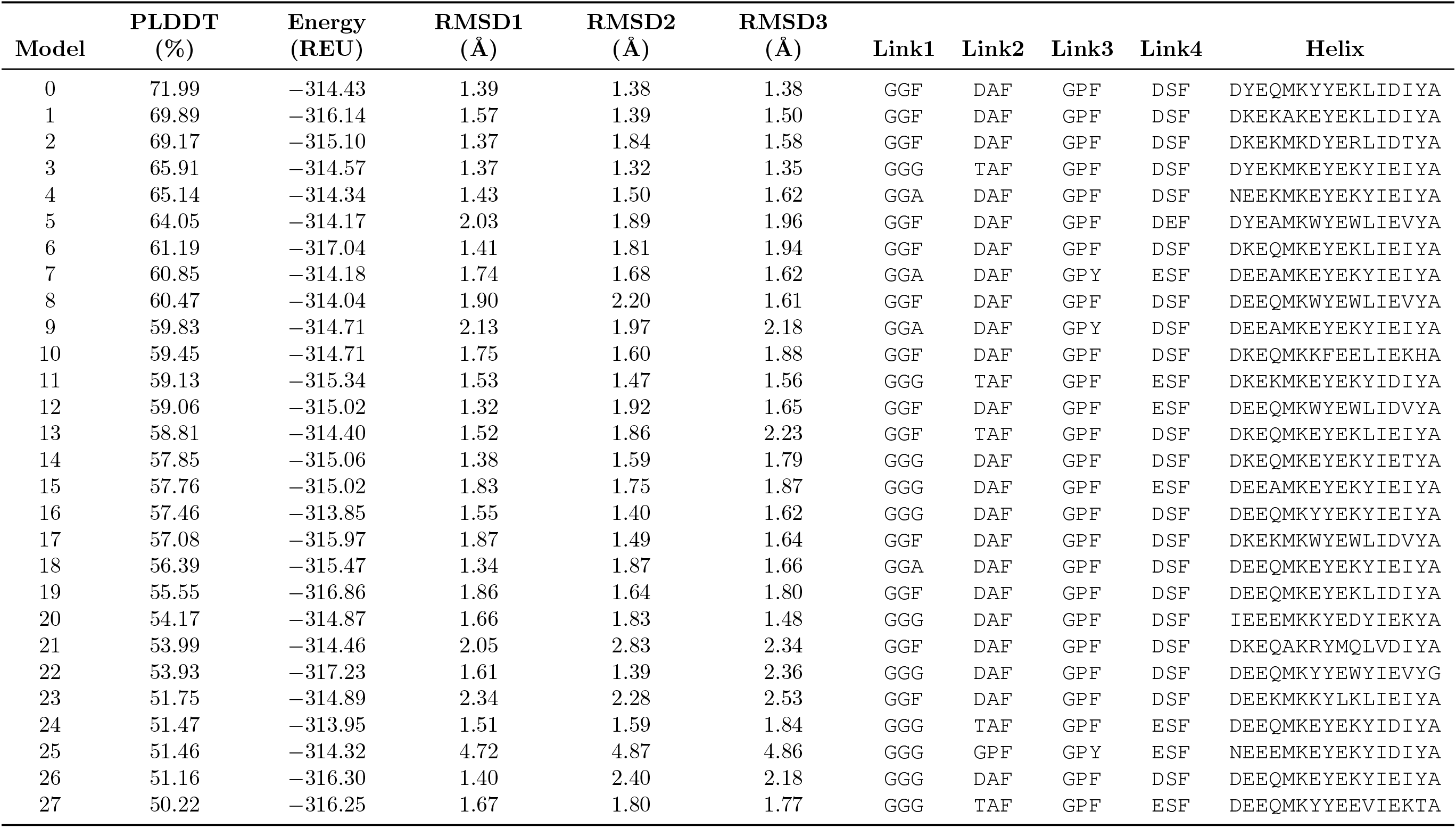
**Models of tau** *β***-helix scaffold.** Attributes of the designed *β*-helix sequences with median (across five AlphaFold^3^ models) of the average per residue confidence (PLDDT) score *>* 50%, and where each of the top three ranking Alphafold predicted structures have RMSD from the Rosetta^1,2^ designed structure *<* 5 Å. The corresponding amino acids for the four linkers as well as the *α*-helix structure for each model are also listed. Models are sorted in order of decreasing values of the median PLDDT score.

